# SARS-CoV-2 spike antigen-specific B cell and antibody responses in pre-vaccination period COVID-19 convalescent males and females with or without post-covid condition

**DOI:** 10.1101/2023.04.13.535896

**Authors:** Marc-André Limoges, Akouavi Julite Irmine Quenum, Mohammad Mobarak H Chowdhury, Fjolla Rexhepi, Mozhdeh Namvarpour, Sara Ali Akbari, Christine Rioux-Perreault, Madhuparna Nandi, Jean-François Lucier, Samuel Lemaire-Paquette, Lakshmanane Premkumar, Yves Durocher, André Cantin, Simon Lévesque, Isabelle J. Dionne, Alfredo Menendez, Subburaj Ilangumaran, Hugues Allard-Chamard, Alain Piché, Sheela Ramanathan

**Affiliations:** Departments of Immunology and Cell Biology; Microbiology and Infectious Diseases and; Department of Biology, Faculty of Science, University of Sherbrooke, Sherbrooke, QC J1K 2R1, Canada; Unité de recherche clinique et épidémiologique, Centre de recherche du CHUS; Department of Microbiology and Immunology, The University of North Carolina at Chapel Hill, Chapel Hill, United States; Medicine, Faculty of Medicine and Health Sciences, Sherbrooke, QC, J1H 5N4, Canada.; Mammalian Cell Expression, Human Health Therapeutics Research Centre, National Research Council Canada, Montreal, QC, Canada; Laboratoire de microbiologie, CIUSSS de l’Estrie – CHUS, Sherbrooke, QC J1H 5N4, Canada; Faculty of Physical Activity Sciences, University of Sherbrooke, Sherbrooke, QC J1K 2R1, Canada.; Research Centre on Aging, Affiliated with CIUSSS de l’Estrie-CHUS, Sherbrooke, QC J1H 4C4, Canada

**Author notes:** Equal first authorship (Marc-André Limoges, Akouavi Julite Irmine Quenum). Shared senior authorship (Hugues Allard-Chamard, Alain Piché, Sheela Ramanathan).

## Abstract

**Background:** Following SARS-CoV-2 infection a significant proportion of convalescent individuals develop the post-COVID condition (PCC) that is characterized by wide spectrum of symptoms encompassing various organs. Even though the underlying pathophysiology of PCC is not known, detection of viral transcripts and antigens in tissues other than lungs raise the possibility that PCC may be a consequence of aberrant immune response to the viral antigens. To test this hypothesis, we evaluated B cell and antibody responses to the SARS-CoV-2 antigens in PCC patients who experienced mild COVID-19 disease during the pre-vaccination period of COVID-19 pandemic.

**Methods:** The study subjects included unvaccinated male and female subjects who developed PCC or not (No-PCC) after clearing RT-PCR confirmed mild COVID-19 infection. SARS-CoV-2 D614G and omicron RBD specific B cell subsets in peripheral circulation were assessed by flow cytometry. IgG, IgG3 and IgA antibody titers toward RBD, spike and nucleocapsid antigens in the plasma were evaluated by ELISA.

**Results:** The frequency of the B cells specific to D614G-RBD were comparable in convalescent groups with and without PCC in both males and females. Notably, in females with PCC, the anti-D614G RBD specific double negative (IgD^-^CD27^-^) B cells showed significant correlation with the number of symptoms at acute of infection. Anti-spike antibody responses were also higher at 3 months post-infection in females who developed PCC, but not in the male PCC group. On the other hand, the male PCC group also showed consistently high anti-RBD IgG responses compared to all other groups.

**Conclusions:** The antibody responses to the spike protein, but not the RBD-specific B cell responses diverge between convalescent males and females, and those who develop PCC or not. Our findings suggest that sex-related factors may also be involved in the development of PCC via modulating antibody responses to the SARS-CoV-2 antigens.

**Short Summary:** Post-COVID Condition (PCC) is lingering illness that afflicts a significant proportion of COVID-19 patients from three months after clearing SARS-CoV-2 infection. Therapy for PCC is only palliative and the underlying disease mechanisms are unclear. The wide spectrum of PCC symptoms that can affect different organs and the detection of viral components in tissues distant from lungs raise the possibility that PCC may be associated with aberrant immune response due to presence of viral antigens. Therefore, we studied B cell and antibody responses to the spike and nucleoprotein antigens in PCC patients who cleared mild SARS-CoV-2 infection during the pre-vaccination COVID-19 pandemic period. We observed divergent patterns of immune reactivity to the spike protein in PCC males and females at different times post-infection, suggesting that the immune responses in PCC may also be influenced by sex-related factors.

## Introduction

SARS and MERS coronaviruses and a few other viral infections are known to cause lingering illnesses (Clark et al., 2015; Hickie et al., 2006; Ong et al., 2004). Among them SARS-CoV-2 is considered unique because of the high proportion of convalescent COVID-19 patients developing the post-covid condition (PCC) (Carfi et al., 2020). PCC encompasses a spectrum of clinical illnesses affecting multiple organs that persist for more than 3 months after the resolution of the initial infection (Nalbandian et al., 2021; Sudre et al., 2021; Sykes et al., 2021). It is estimated that 75-90% of SARS-CoV-2 infected individuals develop a mild COVID-19 disease and 5-50% percent of them develop symptoms of PCC for reasons that still remain unclear (Gallant et al., 2022b; Global Burden of Disease Long et al., 2022). Pre-existing co-morbidities, high level of SARS-CoV-2 viremia, presence of antibodies to type-I IFNs, reactivation of EBV or CMV-specific T cells during acute infection and anti-SARS-CoV-2 antibody signatures are some of the factors suggested to predict PCC in individuals with moderate to severe COVID-19 (Cervia et al., 2022; Peluso and Deeks, 2022; Su et al., 2022). In cohorts of predominantly hospitalized patients, autoantibody titers to type-I IFNs and anti-nuclear antibodies correlate negatively with anti-SARS-CoV-2 antibodies (Su et al., 2022), pointing towards the possibility of deregulated early innate immune response to the virus and/or abnormal activation of atypical memory B cells involved in autoantibody production (Jenks et al., 2018). In PCC associated with mild COVID-19 infection, alterations in certain T cell subsets were observed (Glynne et al., 2022). Peluso et al. (Peluso et al., 2021a) reported a decrease in IFNγ^+^/CD107a^+^ SARS-CoV-2 nucleoprotein-specific CD8^+^ T cells at 9 months post-infection in recovered hospitalized and non-hospitalized COVID-19 patients with PCC, when compared to those without PCC suggesting a possibility of subtle variations in anti-viral immune responses contributing to PCC. These observations also suggest that such altered immune responses towards SARS-CoV-2 antigens in PCC patients may persist long after the resolution of acute COVID-19 disease. To better understand the immune responses towards SARS-CoV-2 antigens in PCC, we focused this study on cryopreserved PBMC and frozen plasma samples collected from unvaccinated individuals who developed PCC following mild, PCR-confirmed COVID-19 disease during the early stages of the pandemic prior to vaccination.

## METHODOLOGY

### Study participants

The Biobanque Québécoise de la COVID-19 (BQC19) is a provincial healthcare initiative undertaken in Quebec, Canada, which collects biological specimens (blood cells and plasma) from individuals with PCR-confirmed SARS-CoV-2 infection and the associated anthropometric and clinical data. Enrolment of adult (>18 years old) participants with different disease severity has been ongoing since 26 March 2020, in several centers throughout the Quebec province. Eligible participants for this study were recruited in the Eastern Townships region of Quebec, between March 2020 and October 2021and were infected before October 2021. Participants were seen at 1, 3, 6, 12, 18 and 24 months after confirmed SARS-CoV-2 infection, and blood samples were obtained during each visit. Acute and persistent symptoms were captured using a 28-symptom questionnaire (**Supplemental document-1**). The questionnaire included details on demographic data including age, sex, weight, height, body mass index (BMI), details of COVID-19 vaccine received and its timing, smoking status, history of hypertension, chronic cardiovascular disease, asthma, other chronic pulmonary diseases, hepatic disease, kidney disease, chronic neurologic disease, active cancer, HIV status, asplenia and use of immunosuppressive drugs. SARS-CoV-2 infection severity was categorized as asymptomatic, mild, moderate and severe according to the WHO definition (WHO, 2021). Peripheral blood mononuclear cells (PBMC) were isolated and preserved in liquid nitrogen and plasma stored at -80°C. This study was approved by the ethic review board of the Centre de Recherche du Centre Hospitalier Universitaire de Sherbrooke (protocol # 2022-4415).

### Detection of RBD specific B cells

To identify RBD-specific B cells, 1-2 × 10^6^ PBMCs were labelled with anti-RBD probes along with a panel of cell surfacemarkers to characterize the different B cell subsets by flow cytometry as previously described (Cohen et al., 2021; Robbiani et al., 2020). Fluorescent RBD probes were made by combining biotinylated strain specific (D614G and Omicron strains) Spike protein RBD tetramers (AcroBiosystems) with fluorescent streptavidin conjugates. The different RBD probes were prepared individually in PBS with a 4:1 tetramer-streptavidin molar ratio. SARS-CoV-2-specific B cells were identified by staining PBMCs with 50ng of labeled RBD tetramer at 4°C for 30 minutes. Each PBMC sample was divided in two halves, one for staining with the D614G RBD-APC/FITC cocktail and the second half with the Omicron RBD-APC/FITC cocktail. The cells were then washed with PBS/FBS-2% and stained for cell surface markers for 30 minutes at 4°C. The cells were washed two times and resuspended in PBS. A viability stain (DRAQ7) was added to each sample 5 minutes before data acquisition using Cytoflex-30 flow cytometer (Beckman Coulter, California, USA). The sources and catalogue numbers of antibodies and reagents are indicated in **Supplemental Table S1**. The data was analyzed using FlowJo software V-10 (BD Biosciences, Mississauga, ON, Canada).

### ELISA for anti-RBD, anti-spike and anti-nucleocapsid antibodies

Anti-RBD specific IgG antibodies were detected by ELISA using a protocol adapted from Moura et al. (Moura et al., 2021). In brief, 96-well high-binding microtiter plates were coated with 50 μL of SMT1-1 spike protein (Colwill et al., 2022), D614G-RBD or omicron RBD antigens at 2 μg/mL in tris-buffered saline (TBS; pH 7.4) (**Supplementary Table S2**). After overnight incubation at 4°C, the plates were washed three times with 200 μL of TBS containing 0.2% Tween 20 (TBST, wash buffer) and blocked with 100 μL of blocking solution (0.1% BSA in TBS containing 0.05% Tween 20) at 4°C overnight. The plates were washed four times with TBST and incubated with plasma samples (**Supplementary Table S3**) TBST-0.1% BSA, in duplicate wells. Samples were incubated for 2 h at 37°C, washed four times with TBST and incubated with the biotinylated anti-human IgG (1:5000), anti-human IgG3 (1:1000) or anti-human IgA (1:10 000) diluted in TBST-0.1% BSA for 1 h at 37°C. Plates were washed and incubated with 50 μL of 3 μg/mL streptavidin–peroxidase diluted in TBST-0.1% BSA for 30 min at 37 °C. After the final washing step (four times), the wells were incubated with the chromogen tetramethylbenzidine (TMB) for 10 min before the reaction was stopped with 50 μL 2N H_2_SO_4_. The optical density at 450 nm (OD450) was measured using SPECTROStar Nano (BMG Labtech, Germany) microplate reader. Plasma samples from 5 convalescent individuals were pooled together to obtain a reference pool. Serial dilutions of the reference pool were used to generate a standard curve with a linear regression equation. The equation was then used to extrapolate the antibody levels in samples taking into account the dilution factor.

Anti-nucleoprotein antibodies were similarly quantified by ELISA using HRP-conjugated secondary antibodies. In brief, 50 μL of NCAP-1 nucleoprotein (Colwill et al., 2022), at 2 μg/mL in TBS was coated in 96-well high-binding microtiter plates overnight at 4°C. The plates were then washed, blocked, incubated with plasma samples, and developed using HRP-conjugated anti-human IgG or anti-human IgA as described above for anti-RBD detection.

### Detection of SARS-CoV-2 mRNA in plasma

Plasma samples were analyzed for the presence of SARS-CoV-2 virus by real-time PCR using the cobas® SARS-CoV-2 Duo test kit using the cobas® 6800 instrument (Roche Molecular Systems, Inc., Branchburg, NJ). Selective amplification of SARS-CoV-2 target nucleic acid sequence in the sample was achieved through the use of a dual target virus specific approach from the highly-conserved regions of SARS-CoV-2 located in the ORF1a and ORF1a/b non-structural regions. The viral load was quantified against a non-SARS-CoV-2 armored RNA quantitation standard (RNA-QS) introduced into each specimen during sample preparation. The RNA-QS also functions as an internal control to monitor the entire sample preparation and PCR amplification process. The system used 0.4 mL of plasma and the viral quantification range of the assay is between 1×10^9^ IU/mL and 100 IU/mL. Below this level of viral load, the SARS-CoV-2 virus can still be detected but not quantified.

### Statistics

GraphPad Prism version 9.5 software was used for statistical analyses and to generate graphics. Statistical comparisons were carried out using the Mann-Whitney’s test. Correlation matrices were calculated using nonparametric Spearman’s correlation. A linear mixed model was used to assess the evolution of antibody response over time. We considered three fixed effects (time, group, and interaction) and a random intercept for repeated measures. Normality of residuals and homoscedasticity (homogeneity of variances) were validated. Results were presented as modelling coefficient (mean differences) and their 95% confidence intervals. Results and figures were obtained using R v.4.1.3. Since these are exploratory results, no multiple testing adjustments were done to obtain *p*-values. A cross-sectional approach was used to analyze the B cell and antibody responses at 1, 3, 6 and 12 months and then a longitudinal study was used in a limited number of participants to assess changes in antibody levels overtime.

## Results

### Increased prevalence of PCC symptoms and their positive correlation to acute COVID-19 disease in females

We undertook an unmatched case control study of comparing convalescent individuals with (PCC) or without (No-PCC) post-COVID condition at 3 months (**Table 1, Fig. 1**). This study also included a small number of uninfected individuals (U, uninfected) who tested negative for the COVID-19 PCR test. In both PCC and No-PCC groups, prior viral infection was confirmed by RT-qPCR and sequential RT-PCR test in nasal swabs was done until a negative test result prior to specimen collection. PCC was defined according to WHO criteria, only when the symptoms persisted for at least 3 months after COVID-19 (WHO, 2021). Participants who received SARS-CoV-2 vaccine post-infection prior to sample collection were excluded from analysis. BMI and frequencies of co-morbidities were comparable between the No-PCC and PCC groups (**Table 1**). Notably, more females were represented in the PCC group than in the No-PCC group in our study cohort (63% versus 38%). In all PCC and No-PCC cases, the acute of disease was mild and none could be categorized as moderate or severe according to WHO criteria. However, the number of the symptoms reported at the acute disease was significantly higher in PCC groups than in No-PCC groups in both males (6 versus 4) and females (7 versus 5) (Mann Whitney test, males- No-PCC versus PCC, p=0.0035; females- No-PCC versus PCC, p<0.0001), in agreement with previous reports (Gallant et al., 2022a; Gallant et al., 2022b; Perlis et al., 2022; Sudre et al., 2021). Notably, in female PCC cases the number of comorbidities positively correlated with age, BMI and the number of PCC symptoms reported at 1-, 3- and 6-months post-infection (**Supplementary Figure S1, supplementary Table S4)**. Moreover, in females the PCC symptoms reported during the acute infection positively correlated with the number of symptoms reported at 1 and 6 months post-COVID, and the symptoms reported at 3 months correlated with those reported at 6 and 12 months in a significant manner. Even though such an impact of acute disease severity with PCC was also observed in males, these correlations were not statistically significant with a few exceptions such as the symptoms reported at the acute and at 1-month post-infection.

**Table 1.** Baseline clinical characteristics of study participants.

| **Variables** | **U**<br>**n = 50** | **No-PCC\***<br>**n = 100** | **PCC\***<br>**n = 87** |
| --- | --- | --- | --- |
| **Male Sex** |  |  |  |
| n (%) | 17 (34.0) | 54 (54.0) | 32 (37.8) |
| **BMI (kg/m<sup>3</sup>) mean (SD)** | 25.9 (4.6) | 27.9 (4.9) | 29.1 (5.6) |
| **Mean age years (SD)** | 40.5 (14.1) | 54.7 (17.2) | 48.2 (13.3) |
| **Age groups** |  |  |  |
| ≤49 years n (%) | 13 (76.5) | 17 (31.5) | 18 (56.2) |
| ≥50 years n (%) | 4 (23.5) | 37 (68.5) | 14 (43.8) |
| **Comorbidities n (%)** |  |  |  |
| Hypertension | 2 (11.7) | 19 (35.2) | 5 (15.6) |
| Diabetes | 0 (0) | 7 (12.9) | 2 (6.3) |
| Asthma | 1 (5.8) | 2 (3.7) | 1 (9.0) |
| Autoimmune diseases | 4 (23.5) | 6 (11.1) | 3 (9.3) |
| **Number of symptoms at the acute phase median value (range)** |  | 4 (1-13) | 6 (0-13)<sup>@</sup> |
| **Female Sex** |  |  |  |
| n (%) | 33 (66) | 46 (46) | 55 (63.2) |
| **BMI (kg/m<sup>3</sup>) mean (SD)** | 26.5 (5.2) | 27.1 (7.6) | 27.1 (5.6) |
| **Mean age years (SD)** | 38.1 (14.8) | 46.7 (17.6) | 48.5 (16.4) |
| **Age groups** |  |  |  |
| ≤49 years n (%) | 27 (81.8) | 24 (52.2) | 28 (50.9) |
| ≥50 years n (%) | 7 (21.2) | 22 (47.8) | 27 (49.1) |
| **Comorbidities n (%)** |  |  |  |
| Hypertension | 2 (6.1) | 3 (6.5) | 14 (25.5) |
| Diabetes | 0 (0) | 0 (0) | 2 (3.6) |
| Asthma | 1 (3.0) | 3 (6.5) | 1 (9.0) |
| Autoimmune diseases | 4 (23.5) | 6 (11.1) | 8 (14.5) |
| **Number of symptoms at the acute phase median value (range)** |  | 5 (1-8) | 7 (2-12)<sup>§</sup> |

\* According to WHO classification (WHO, 2021).
U- uninfected; No-PCC convalescent individuals without PCC; PCC- convalescent individuals with PCC; PCC= post-covid condition; SD, standard deviation.
<sup>@</sup> Comparison with No-PCC group p=0.0035; Mann Whitney test.
<sup>§</sup> Comparison with No-PCC group p<0.0001; Mann Whitney test.

**Figure 1:**
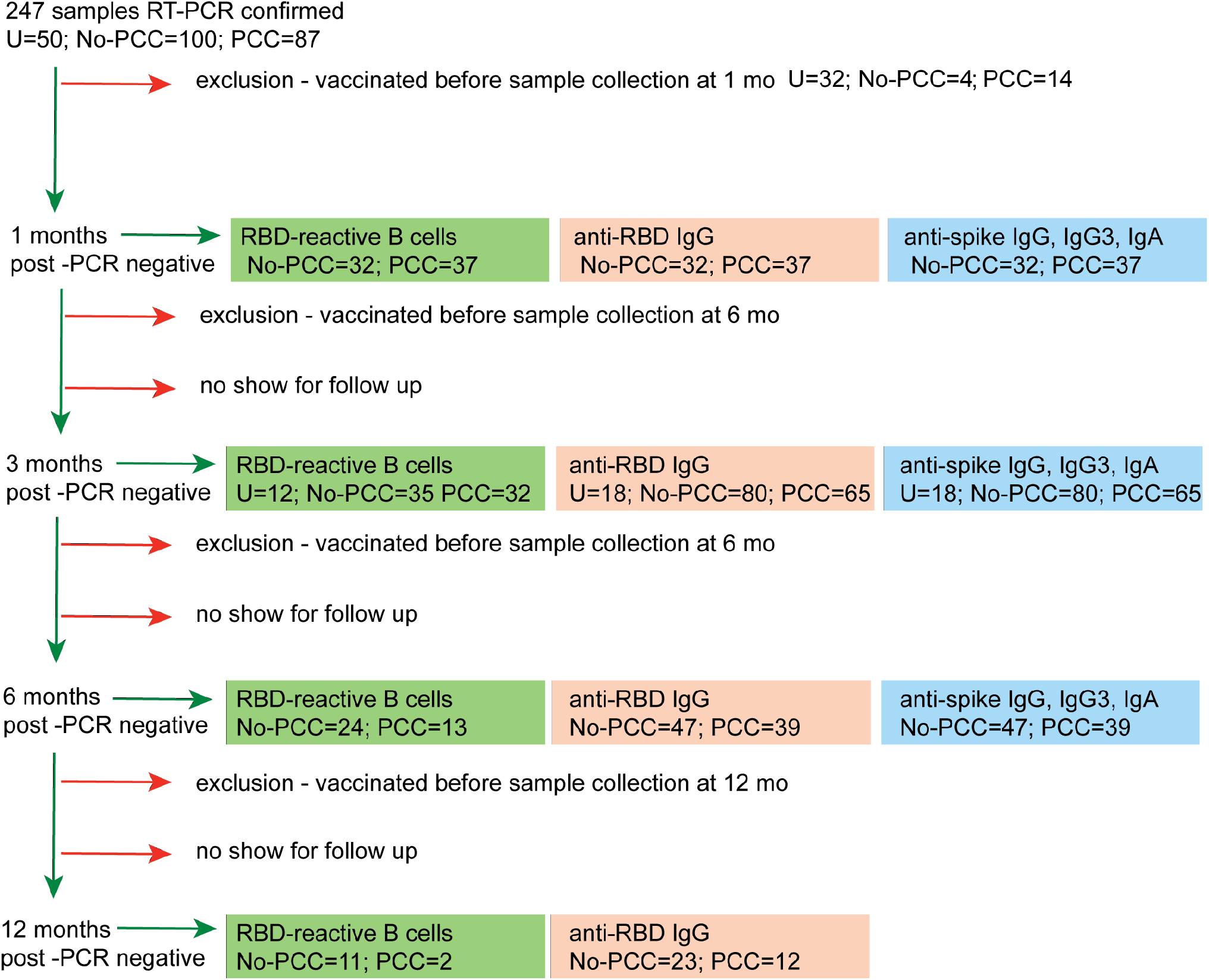
Workflow of sequential immune reactivity analysis of convalescent COVID-19 subjects with or without PCC and sample size distribution. All samples were obtained from the BQC-19 biobank for COVID-19. At recruitment stage, negative COVID-19 status was confirmed by RT-PCR. Sequential RT-PCR tests for SARS-CoV-2 was carried out in COVID-19 positive individuals to confirm virus clearance before specimen collection. PCC, convalescent individuals with persistent symptoms at 3 months post-COVID-19. NO-PCC, no PCC symptoms after acute infection. Plasma was available for experimentation from all PCC or No-PCC individuals at 3 months post-infection. PBMCs were available for experimentation for a subset of individuals. Some individuals had samples available at 1-month post infection. Significant proportion of individuals dropped out at 6- and 12- months during follow up. Individuals who had received SARS-CoV-2 vaccine at any time before sample collection at 1, 3 or 6 months were excluded from the analyses for these time points. Healthy PCR negative individuals with no history of COVID-19 symptoms constituted the uninfected controls (U). Samples from uninfected controls were collected at one time point only.

### RBD-specific memory B cell frequencies are comparable between PCC and No-PCC groups at three months post-infection

During the early stages of viral infections, antigen-specific cognate B cells may directly differentiate into antibody producing cells in the extra-germinal space (Beckers et al., 2023; Ruschil et al., 2020). As the antiviral immune response progresses with the activation of T cells germinal centers are formed, somatic hypermutation occurs and long-lasting memory B cells are generated (Ruschil et al., 2020; Sanz et al., 2019; Sokal et al., 2021). The loss of surface IgD and the expression of CD27 are hallmarks of B cells undergoing maturation in germinal centers (Klein et al., 1998). In addition, IgD^-^ CD27^-^ (double negative, DN) B cells are also reported to be enriched during extrafollicular B cell maturation associated with certain inflammatory conditions and infections including COVID-19 and HIV (Kaneko et al., 2020; Krause et al., 2022; Ruschil et al., 2020; Sanz et al., 2019; Sokal et al., 2021). SARS-CoV-2 infection has been reported to induce viral antigen specific memory B cells within germinal center follicles as well as outside the follicles (Kaneko et al., 2020; Sokal et al., 2021; Woodruff et al., 2020). Therefore, we first assessed whether PCC was associated with altered SARS-CoV-2-specific memory B cells, using RBD region as a surrogate antigen to detect such cells in PBMC.

We assessed the B cell reactivity of convalescent PCC and No-PCC samples as well as SARS-CoV-2 PCR-negative, unvaccinated control samples toward the RBD epitope (referred to as D614G RBD) (https://cov-lineages.org/resources/pangolin.html). Nonetheless, we also assessed B cell reactivity toward the omicron-RBD to assess potential cross reactivity. The gating strategy for identifying RBD specific B cells is shown in **Supplementary Fig. S2**. The staining pattern of CD27 and IgD on CD19^+^CD20^+^ B lymphocytes identified naïve (naïve, IgD^+^CD27^-^), unswitched memory (USM, IgD^+^CD27^+^), switched memory (SM, IgD^-^CD27^+^) and double negative (DN, IgD^-^CD27^-^) B cells. The reactivity of these B cell subsets to D614G-RBD and omicron-RBD was evaluated using epitope-specific tetramers. The frequency of plasmablasts (CD19^+^CD20^-^) in PBMC were very low, hence these cells were not analyzed for RBD binding.

PCC and No-PCC groups displayed a significantly elevated proportion of D614G-RBD reactive B cells within total CD19^+^ and CD19^+^CD20^+^ (excludes plasmablasts) B cell pools when compared to uninfected controls (**Fig. 2a,b**; left panels). The increased frequency of RBD specific B cells mainly resulted from class-switched (IgD^-^) B cells (**Fig. 2c**; left panel), whereas IgD^+^ B cells (constitutes unswitched and naïve) showed very low reactivity (**Fig. 2d**; left panel). Comparison of the frequencies of D614G-RBD specific B cell subsets between males and females showed no significant differences between the two convalescent groups (**Supplementary Fig. S3**). Hence, further analyses on RBD reactivities were carried out on pooled data without segregating them by sex. Naïve B cells lose surface IgD expression following antigen mediated activation and class switching. IgD^-^CD27^+^ SM B cells that undergo T cell dependent class switching in germinal centers, showed significantly increased frequencies of D614G-RBD reactive cells in the two convalescent groups compared to uninfected controls (**Fig. 2e**; left panel). IgD^-^CD27^-^ DN B cells, which predominantly originate from extrafollicular class switching, also harbored significantly more D614G-RBD specific cells in both PCC and No-PCC convalescent groups compared to uninfected controls (**Fig. 2f**; left panel).

**Figure 2.**
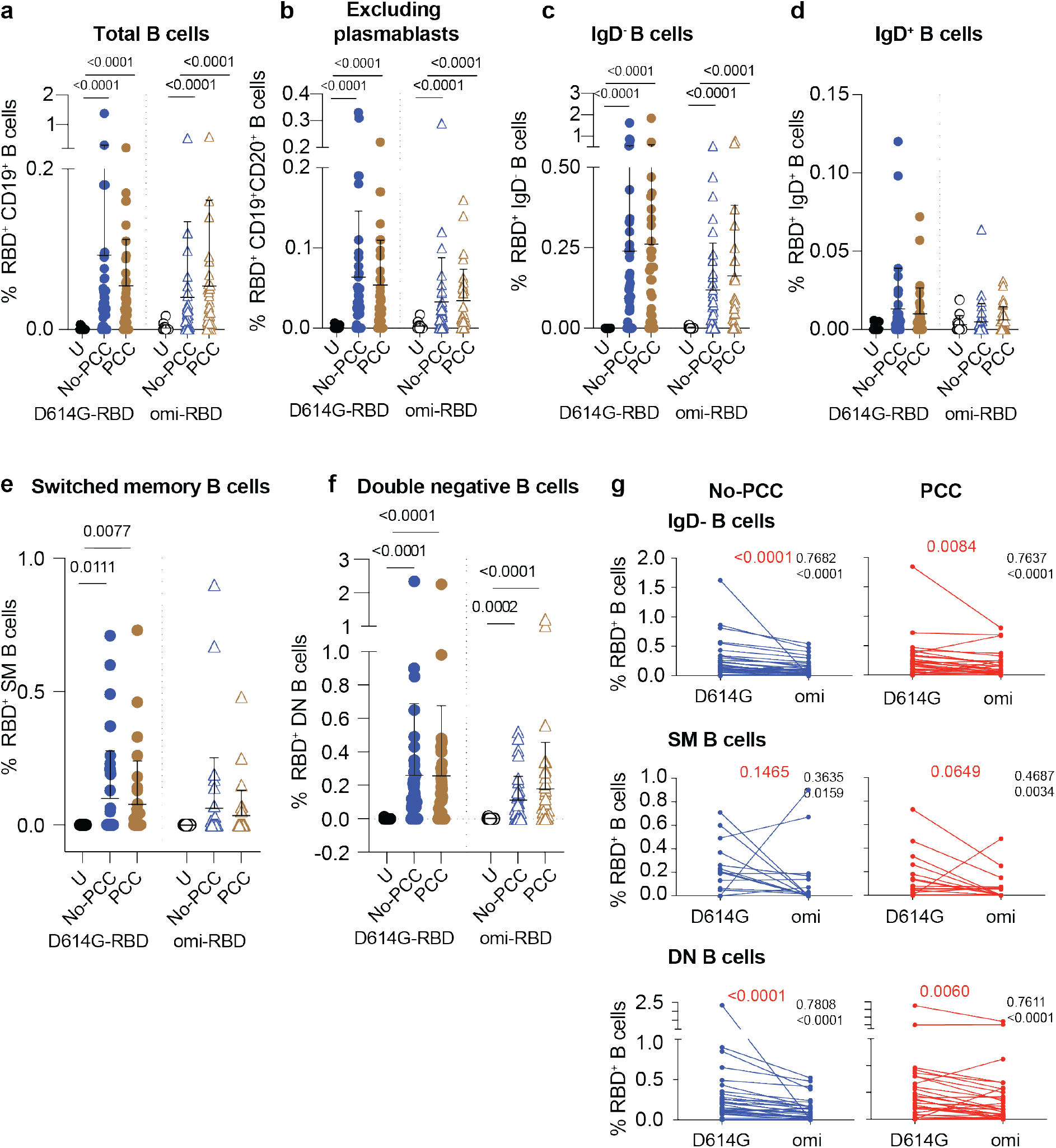
Frequencies of RBD specific B cell subsets at 3 months after mild SARS-CoV-2 infection. PBMCs were labelled with D614G-RBD or omicron-RBD and specific B cell markers, and B cell subpopulations were gated as shown in Supplementary Figure S2. (**a-f**) Frequencies of D614G-RBD and omicron-RBD specific B cells in total CD19^+^ B cells (**a**). CD19^+^CD20^+^ (plasmablasts excluded) B cells (**b**). CD19^+^CD20^+^IgD^-^ B cells (**c**). CD19^+^CD20^+^IgD^-^ B cells (**d**). CD19^+^CD20^+^CD27^+^IgD^-^ switched memory (SM) B cells (**e**). CD19^+^CD20^+^CD27^-^IgD^-^ double negative (DN) B cells (**f**). The groups were compared by Mann Whitney’s test and significance values are indicated. Sample numbers: Uninfected (U); convalescent individuals without PCC (No-PCC); convalescent individuals with PCC (PCC). (**g**) Correlation between anti-D614G-RBD and omicron-RBD specific B cell frequencies in No-PCC and PCC groups. The values in black are *ρ* value for Spearman coefficient and the corresponding *p* value. The values in red represent the *p* values for comparisons between D614G and omicron-RBD specificities by Wilcoxon matched-pairs signed rank test.

Evaluation of B cell reactivity to omicron-RBD also showed significantly increased frequencies within total CD19^+^ and CD19^+^CD20^+^ B cell pools and in the IgD^-^ subset in No-PCC and PCC groups compared to uninfected controls (**Fig. 2a-c**; right panels). Reflecting the specificity of SM B cells to the eliciting antigen-D614G-RBD, the omicron specificity in SM B cells in the convalescent groups was not significantly different from the uninfected group (**Fig. 2e**; right panel). However, omicron-RBD specific B cell frequencies were significantly higher in the DN subset of both PCC and No-PCC convalescent groups than in uninfected controls (**Fig. 2f**; right panel). The frequencies of omicron specific B cells were markedly lower than that of D614G-RBD specific B cells in IgD^-^ and DN B cell subsets (**Fig. 2c, f**). However, the anti-D614G and anti-omicron-RBD responses showed significant correlation in these B cells subsets in both No-PCC and PCC groups (**Fig. 2g**), suggesting potential cross-reactivity between D614G and omicron-RBD that may confer partial protection to the omicron variant.

### Distinct anti-RBD and anti-spike antibody responses in PCC and No-PCC groups at three months post-infection

Parallel analysis of antibody titers to SARS-CoV-2 antigens revealed significantly elevated anti-D614G-RBD IgG antibody titers in the plasma samples obtained at three months post-infection in both PCC and No-PCC groups compared to uninfected controls (**Fig. 3a**). Notably, the PCC group showed significantly higher levels of anti-D614G-RBD IgG response than the No-PCC group (**Fig. 3a**), even though their D614G-RBD specific B cell frequencies were comparable within IgD^-^, DN and SM B cell subsets (**Fig. 2**). On the other hand, IgG response to omicron-RBD was significantly increased in No-PCC and PCC groups when compared to uninfected controls but was not different between the two convalescent groups, suggesting a differential response to omicron epitopes in PCC (**Supplementary Fig. S4**). Segregation of the serology data by sex showed that the significantly elevated anti-D614G-RBD IgG responses of PCC group compared to No-PCC group was recapitulated only in males with PCC but not in females (**Fig. 3b**). Moreover, within the PCC group males displayed significantly higher anti-D614G-RBD IgG levels than females (**Fig. 3b**). However, analysis of the antibody response against the whole spike protein showed a different pattern in males and females. Anti-spike IgG, IgG3 and IgA levels were significantly higher in females with PCC when compared to No-PCC females, whereas these responses were comparable between PCC and No-PCC males (**Fig. 3c, d, e**). Within the No-PCC group, anti-spike IgG and IgA levels were markedly higher in males than in females, but this was not seen within the PCC group (**Fig. 3c, d, e**). In contrast to anti-spike antibody responses, anti-nucleocapsid IgG responses were similarly elevated in both PCC and No-PCC groups when compared to uninfected controls (**Fig. 3f**). However, there were no differences in anti-nucleocapsid IgG and IgA responses between males and females in both groups (**Fig. 3g**). Overall, the anti-D614G-RBD IgG levels were significantly higher in the PCC group than in No-PCC groups among males, whereas anti-spike IgG, IgG3 and IgA levels were elevated in PCC than in No-PCC group among females at three months post-infection.

**Figure 3:**
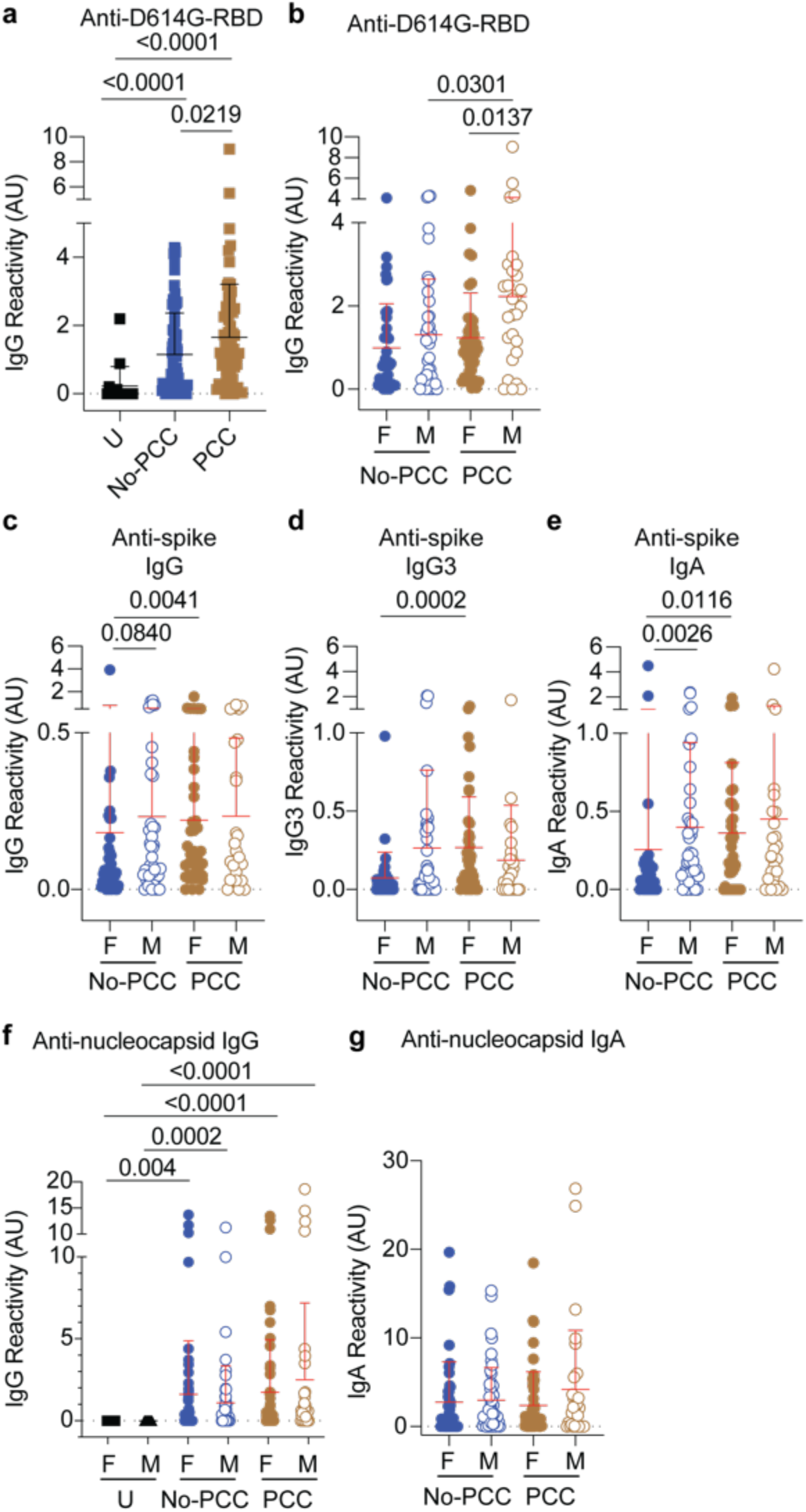
Anti-RBD, spike and nucleocapsid antibody responses in No-PCC and PCC groups at 3 months post-infection. Plasma samples were collected during routine clinical visit at 3 months post PCR-positive diagnosis. Total IgG, IgG3 and IgA responses in uninfected (U) controls and convalescent PCC and No-PCC groups were determined by ELISA. (**a**) anti-D614G responses in pooled (males and females) samples. (**b**) anti-D614G-RBD IgG responses in males and females within PCC and No-PCC groups. (**c-e**) Anti-spike IgG, IgG3 and IgA responses in males and females with or without PCC. (**f-g**) Anti-nucleocapsid IgG and IgA responses in males and females within U, PCC and No-PCC groups. The groups were compared by Mann Whitney’s test.

### Pattern of immune responses at three months distinguish male and female convalescent groups with and without PCC

Next, we determined how the symptoms at acute infection correlated with the B cell reactivities and antibody responses at three months post infection in convalescent PCC and No-PCC groups in males and females using correlation matrices. The Spearman correlation and the corresponding significance values depicted in **Fig. 4 (Supplementary Table S5)** showed a significant positive correlation between anti-spike IgG and anti-D614G-RBD IgG antibodies in all groups, as expected (**Fig. 4**, orange asterisks). Similarly, anti-D614G-RBD specific DN B cells showed a significant positive correlation with D614G-RBD specific CD19^+^CD20^+^ B cells and the class-switched IgD^-^ subset in all the groups (**Fig. 4**, yellow asterisks), indicating that our observations on the D614G-RBD specific DN B cells reflect the B cell responses to the RBD domain in these individuals as expected. Curiously, the IgG, IgG3 and IgA response to spike protein showed significant correlation to D614G-RBD specific CD19^+^, CD19^+^CD20^+^ and IgD^-^ B cell responses only in the convalescent males who do not develop PCC at 3 months post infection expected (**Fig. 4b**, blue asterisks). In all the other groups (males and females with PCC and females without PCC), the antibody responses showed no specific correlation to D614G-RBD specific B cell responses, despite the presence of detectable levels of circulating antibodies to spike and D614G-RBD (**Fig. 4a, c, d**; **Supplementary Table S5, Fig. 3**). Notably, the number of symptoms at the acute infection did not correlate with any B cell or Ab parameters in male or female PCC cohorts except with anti-D614G-RBD specific IgD^-^ B cells in females with PCC (**Fig. 4c**, red asterisk). Female PCC cases also showed a strong positive correlation between CD19^+^ and SM B cells that was not observed in other groups (**Fig. 4c**, green asterisks). Collectively, these observations indicate that the convalescent males show a coordinated immune response that appear to be deregulated by PCC in males.

**Figure 4.**
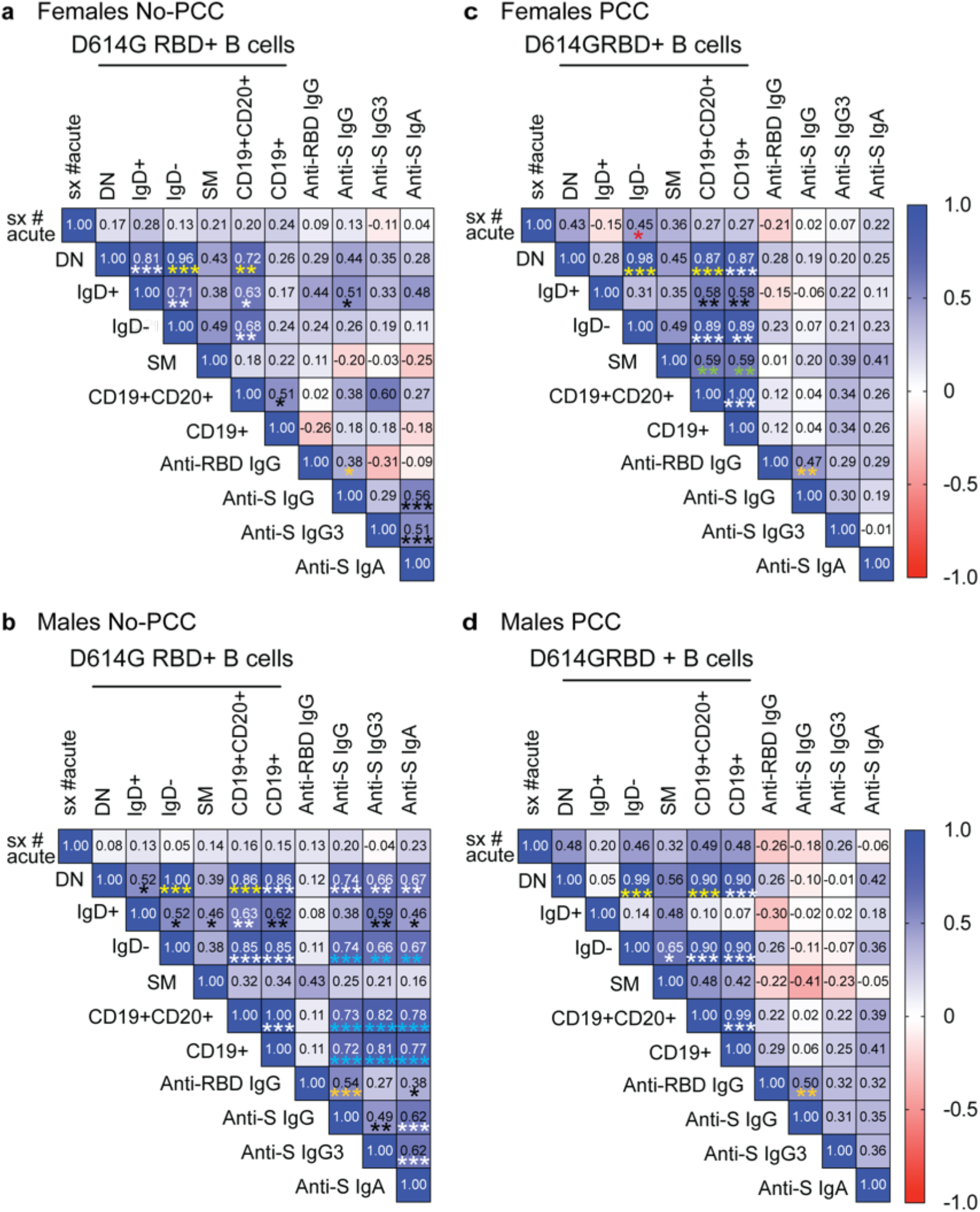
Correlation between SARS-CoV-2 Spike protein-specific B cell reactivities and antibody responses in PCC and No-PCC groups among males and females. Correlation matrices were generated for the indicated parameters for the convalescent groups with or without PCC. Nonparametric Spearman’s correlation coefficient values (numbers) and their significance (asterisks) are indicated. * *p*<0.05, ** *p*<0.01, *** *p*<0.001. Asterisks are color coded to as visual aids to denote specific comparisons described in the text. The white and black numbers and asterisks are only used to contrast with the background color. The number of samples and exact *p* values are shown in the **Supplementary Table S5**.

To determine how the B cell reactivities and antibody responses towards SARS-CoV-2 antigens during the early convalescent period correlated with progression towards PCC, we evaluated these immune parameters at 1-month post-infection. The proportion of D614G-RBD specific B cells were comparable between the two convalescent groups in females and males at 1-month post-infection as at three months (**Figs. 5a, 2a and Supplementary Fig. S3**). D614G-RBD reactive IgD-B cells, anti-D614G-RBD and anti-spike IgG levels tended to be higher in males with PCC when compared to males without PCC at 1-month post-infection (**Fig. 5a, b**) even though these parameters were comparable at 3-months post-infection among males (**Fig. 3c, Supplementary Fig. S3**). An inverse trend was noted for female PCC and No-PCC groups, with significantly higher anti-spike IgG response in the PCC group at 3 months but not at 1-month post-infection. Antibody responses to nucleocapsid antigens were comparable between the PCC and No-PCC group among males and females at 1-month post-infection as at 3-months (**Fig. 5d and 3g,f**). Our study cohort included a few individuals who got infected after vaccination among males and females within the No-PCC group and females within the PCC group. These vaccinated individuals exhibited higher anti-D614G RBD and anti-spike IgG responses but reduced antibody responses to nucleocapsid at 1-month post-infection (**Fig. 5a-d**), however, these sample numbers are too low to permit meaningful comparisons.

**Fig 5.**
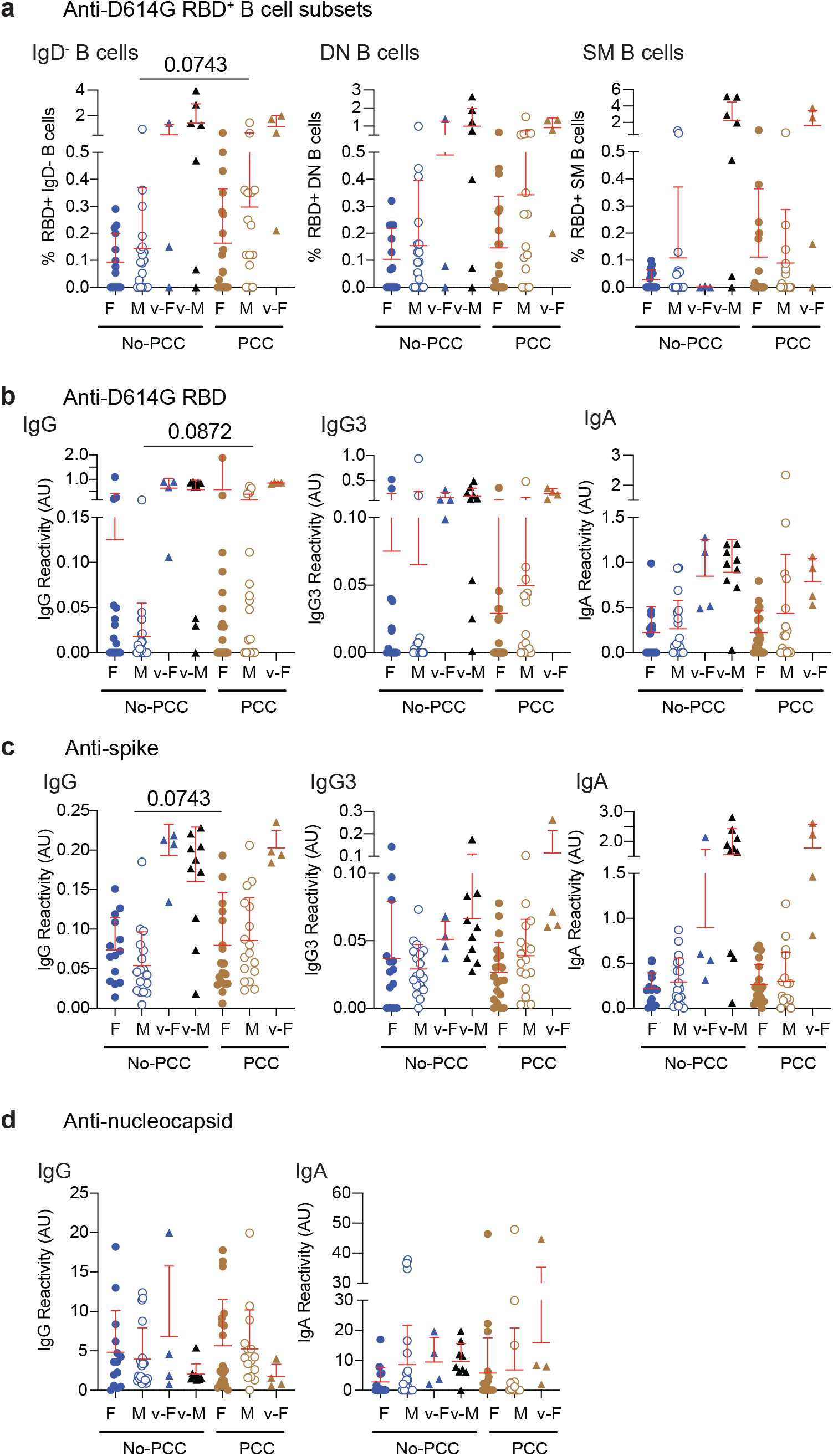
Anti-RBD, spike and nucleocapsid specific responses in samples obtained at 1-month post-infection. Plasma samples were collected during routine clinical visit at 1 month after PCR-positive diagnosis. (**a**) D614GRBD reactive B cell subsets. (**b**) anti-D614G-RBD (**c**) anti-spike and (**d**) anti-nucleocapsid antibody responses were determined in females and males with or without PCC. v-M and v-F denote vaccinated individuals within the indicated groups. The groups were compared by Mann Whitney’s test.

PCA analyses of the data obtained at one- and three-months post-infection indicated clear differences between the two timepoints but did not distinguish between convalescent individuals with and without PCC (**Supplementary Fig S5**). Nonetheless, we examined the concordance between the different antibody responses in the four study cohorts at 1-month post-infection. The anti-D614G-RBD and anti-spike antibody responses were comparably coordinated in No-PCC groups among males and females (**Fig 6a,b; Supplementary Table S6**). However, in the females who developed PCC, anti-D614G-RBD and anti-spike antibody responses showed a high degree of correlation with anti-nucleocapsid responses (**Fig 6c**, yellow asterisks), whereas these correlations were mostly non-significant or weak in the male PCC group. In all the four study groups, the anti-D614G-RBD levels did reveal any meaningful correlation with the D614G-RBD-reactive B cell subsets (**Supplementary Fig S6**). In both males and females with PCC, the symptoms at the acute significantly correlated with anti-D614G-RBD IgG (Supplementary **Fig 6c,d**), although they did not correlate at 3 months post-infection (**Fig. 4c,d**). These observations suggest that the IgG response to D614G-RBD at 1-month post-infection appears to be a notable hallmark to predict PCC in males and females.

**Fig 6.**
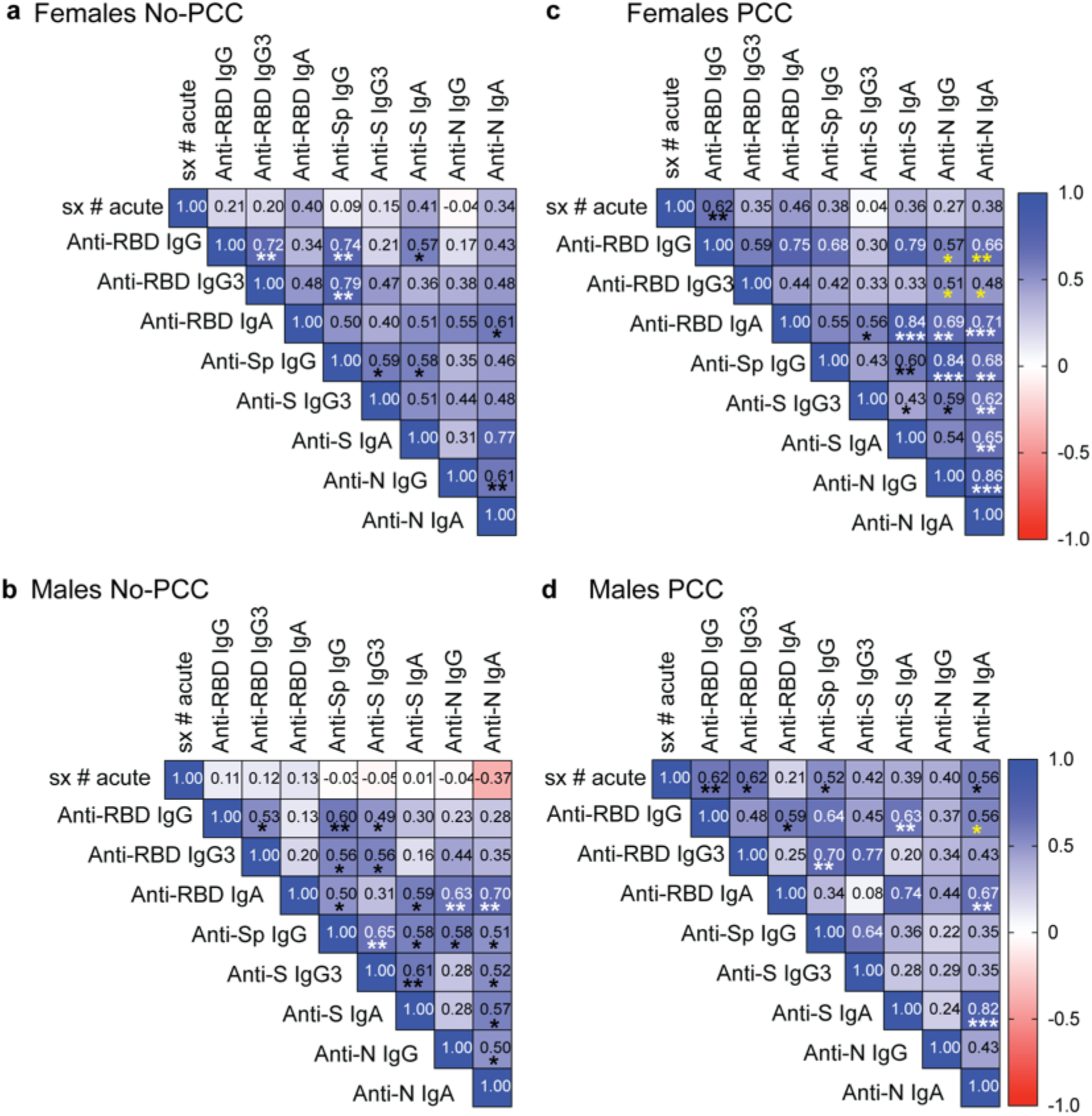
Correlation matrix for anti-RBD, spike and nucleocapsid antibody responses in PCC and No-PCC groups at 1-month post-infection. Correlation matrix for antibody responses at 1-month post infection in (**a**) females No-PCC, (**b**) males No-PCC, (**c**) females PCC and (**d**) males PCC groups. RBD refers to D614G-RBD. The numbers in the squares indicate the Spearman coefficient value. The *p* values are indicated in the figure as asterisks (* *p*<0.05, ** *p*<0.01, *** *p*<0.001) and are given in **Supplementary Table S6.** Due to the limited quantity of samples in which B cell reactivities were measured, these parameters were not included in the correlation matrix and are given in **Supplementary Fig. S6**.

### D614G-RBD specific IgG levels persists at 6 months post infection in convalescent males with PCC group

As the strength of adaptive immune response to SARS-CoV-2 is proportional to severity of infection, attrition of SARS-CoV-2 specific antibodies and B cells is observed in mildly symptomatic infections over time (Beaudoin-Bussieres et al., 2020; Crawford et al., 2021; Dan et al., 2021; Figueiredo-Campos et al., 2020; Gaebler et al., 2021; Robbiani et al., 2020; Rodda et al., 2021; Wheatley et al., 2021). Sequential plasma and PBMC samples obtained from the same individuals at one-, three-, six- and twelve-months post-infection and were analyzed for anti-D614G-RBD responses. Only individuals who had not been vaccinated at the time of sample collection at 6 months post-COVID-19 diagnosis were included in the analyses. Anti-D614G-RBD antibody response was barely detectable in male and female convalescent groups without PCC, whereas males in the PCC group showed persistently high levels of anti-D614G-RBD IgG levels at 6 months post-infection (**Fig. 7a**). Evaluation of the longevity of these antibody responses at 1, 3 and 6 months showed a rise in anti-D614G-RBD IgG levels at 3 months posit-infection that returned to baseline levels on No-PCC males and females, whereas it remained elevated in a few females and in a significant proportion of male PCC cases, which is reflected in a significant interaction value between the 3 and 6 months timepoints (**Fig. 7b, c**). Interestingly, anti-spike IgG, IgG3 and IgA titers did not show appreciable decline at 6 months in both No-PCC and PCC groups among males and females (**Supplementary Fig S7**). Comparison of the pattern of antibody responses to spike and RBD at 1-, 3- and 6-months post infection, also revealed a better coordinated immune response in males without PCC over time (**Supplementary Fig. S8, Supplementary Table S7**). In females without PCC, the antibody responses at 1 and 3 months appear to show some correlation, but not at later time points. On the other hand, in males and females with PCC, these obvious patterns were not evident. Taken together, these observations point towards the possibility that the immune responses generated at early time points (1 to 3 months) after the infection may be associated with PCC.

**Figure 7.**
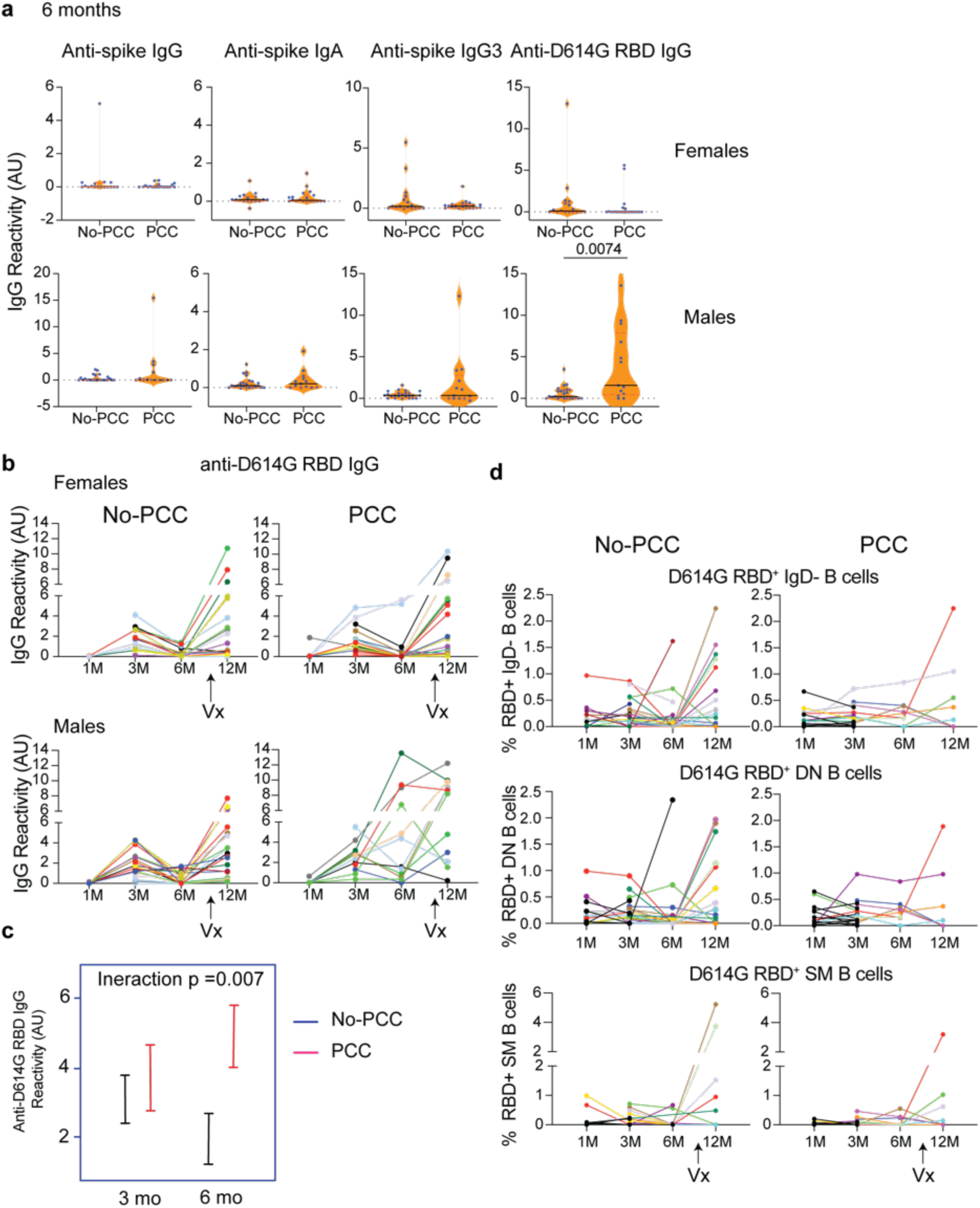
Persistent D614G-RBD specific IgG antibodies at 6 months in the male PCC group. Plasma samples acquired at 1-, 3-, 6- and 12-months post-SARS-CoV-2 PCR positivity were analyzed for anti-D614G-RBD specific IgG antibodies as described in Figure 3. (**a**) Anti-spike IgG, IgG3 and IgA antibodies and anti-D614G-RBD specific IgG antibodies at 6 months post-infection in non-vaccinated No-PCC and PCC groups among females and males. (**b**) evolution of anti-RBD IgG antibody from 1-12 months in females and males without or with PCC. (**c**) Comparison of the changes in the anti-RBD IgG between convalescent males without (blue) or with PCC (red) from 3 to 6 months. (**d**) Changes in RBD specific B cells. Statistical analysis was carried out by Mann-Whitney’s test (**a**) and by linear mixed model (**c**).

At 12 months, most individuals included in this study had received at least 1 dose of vaccine (Pfizer, Moderna or Comirty). Vaccination in general induced a strong anti-D614G-RBD IgG response in all groups to a comparable level, although significant numbers of non-responders were evident in all groups (**Fig. 7b**). This increase in IgG response was associated with a trend towards an increase in the proportion of D614G-RBD reactive IgD^-^, DN and SM B cells that was more apparent in No-PCC cases than PCC cases, which were limited by small sample size for statistical comparisons (**Fig. 6d, Supplementary Fig. S9**).

## Discussion

To our knowledge this is the first follow-up study on convalescent COVID-19 subjects who recovered from the disease but developed PCC that analyzes immune responses towards SARS-CoV-2 antigens at different time points post-infection. The results presented here indicate that individuals with PCC have persistent and/or altered B cell and antibody responses towards the spike protein, and not nucleocapsid protein. As the study included only the subjects with mild COVID-19 disease, the overall antibody titers were low in both PCC and control No-PCC convalescent groups in agreement with other reports (Beaudoin-Bussieres et al., 2020; Crawford et al., 2021; Dan et al., 2021; Gaebler et al., 2021; Rodda et al., 2021; Wheatley et al., 2021). However, in both male and female PCC subjects the number of symptoms at the acute disease correlated with anti-RBD IgG antibody level at 3 months post-infection (**Figs. 4**). Whereas the anti-spike antibody responses are higher in convalescent females who develop PCC, males with PCC predominantly display higher anti-RBD IgG responses (**Figs. 3, 5, 6**). As early as one-month post-infection, the number of symptoms at the acute disease correlated with anti-RBD IgG antibody level in both males and females (**Fig. 6**). Anti-RBD B cell responses were comparable between convalescent males and females and between those with and without PCC. However, differences were observed at the level of circulating antibodies to spike antigen. Given the differences in the severity of the symptoms at the acute disease and the differential susceptibility of males and females to PCC, we analyzed the immune responses separately in the two sexes.

### Differences in the antibody responses between convalescent males and females

The immune responses to the spike antigen seem to differ between males and females after mild COVID-19. At 3 months post-infection, the magnitude of anti-spike antibody responses is lower in the No-PCC convalescent females when compared to No-PCC convalescent males (**Fig. 3**) even though they were comparable at one-month post-infection (**Fig. 5**). The anti-RBD specific B cells, used as a surrogate for B cell responses to SARS-CoV-2 antigens, were also comparable between the sexes irrespective of their PCC status at 1, 3 and 6 months post-infection (**Figs. 2, 5, 7**), supporting the observations that the sex-dependent influences are manifested following the generation of the antigen-specific B cells (Paavonen et al., 1981; Sthoeger et al., 1988).

Anti-viral immune responses are reported to be reduced in amplitude in females when compared to males during the early stages of COVID-19 pandemic and for other infections at the mucosal surfaces (Falagas et al., 2007; Grzelak et al., 2021; Mitchell et al., 1992; Ozgocer et al., 2022; Zhai et al., 2022). Susceptibility to SARS, MERS and SARS-CoV-2 infections and viremia tends to be higher in males than in females (Peckham et al., 2020). In contrast to SARS infections, females were more susceptible during 1957 and 2009 influenza pandemics (Kilbourne, 2006; Morgan and Klein, 2019; Serfling et al., 1967). Innate immune responses to viral infections have been reported to be higher in females (Kovats, 2015; Regis et al., 2021), which can explain the increased responsiveness of females to vaccines (Fischinger et al., 2019). COVID-19 vaccines elicited higher levels of IgG antibodies in females than in males, probably reflecting the heightened innate immune responses in females (Demonbreun et al., 2021; Ebinger et al., 2022). The increased responsiveness of female biological sex when compared to male biological sex has been observed across various vaccination protocols even in younger age groups (Ohm et al., 2022; Voysey et al., 2016; Webster et al., 2022). Similarly, females are more prone to develop autoimmunity (reviewed in (Klein and Flanagan, 2016)). In the light of the above reports, our results showing that the anti-spike responses are lower in convalescent females suggest that the sex-based differences in susceptibility to infections, immune response to vaccines and the propensity to develop immune-related pathologies (PCC here and autoimmunity in discussion) are likely modulated in context dependent manner such as the nature of antigen exposure (vaccine versus infection), the nature of the antigen encountered and the pathogen in question.

Another notable observation of our study is the positive correlation between anti-spike B cell and antibody responses at 1 month in both females and males irrespective of whether they progress toward PCC or not (**Fig. 6**). In mild COVID-19 disease, coordinated innate, T and B cell responses at 3 weeks post infection has been observed (Capelle et al., 2022). However, by 3 months post-infection, this correlation becomes stronger in the males who do not develop PCC whereas in other groups studied such correlation is restricted to the D614G-RBD specific B cell subsets (**Fig. 4**). Anti-spike IgG and anti-RBD antibody responses showed similar positive correlations at 1- and 3-months post-infection in males and females who did not develop PCC. At 12 months post-infection, where most of the samples were from vaccinated individuals, the anti-RBD B and antibody responses were highly coordinated (**Supplementary Fig. S8**). Thus, the differences seen following natural infection in convalescent males and females with PCC were probably overshadowed/minimized by the effect of adjuvants in the vaccine. Omics based systems biology approach indicated that males who recovered from COVID-19 showed a better coordinated innate, B cell and antibody responses to subsequent influenza vaccination (Sparks et al., 2023). While the history of infections and influenza vaccinations are not known in the participants involved in this study, nevertheless it suggests that immune responses are continually being shaped by past infections. It is possible that SARS infections shape the immune responses in males and females in a distinct manner such that immune responses to a heterologous antigen, such as influenza vaccine studied by Sparks et al., are altered (Sparks et al., 2023). Immunodominance hierarchies have been observed to be have a sex/gender bias in T cell responses (Mifsud et al., 2008; Zhang et al., 2008). Thus, it is not unlikely that each infection displays a distinct sex bias that influences the outcome of subsequent infections/vaccinations. As data comparing the immune responses of post-puberty males and females to other infections are unavailable to our knowledge, it is difficult to determine whether our observations on convalescent COVID-19 males and females are unique to SARS-CoV-2 infection or can be generalized to other infections. While trained innate immune response following SARS-CoV-2 infection in males appear to correlate with the outcome of influenza vaccination (Sparks et al., 2023), it raises the possibility that immune responses in males are continuously shaped by the previous infections while they may be subject to modulation by sex-intrinsic factors in females.

### B cell responses

In most of the convalescent individuals with or without PCC, we observed that the RBD specific B cells were predominant in circulation, although there was no significant correlation between the RBD specific B cell subsets and anti-RBD specific antibody levels, as reported in SARS-CoV-2 vaccinated individuals (Nayrac et al., 2022). While IgD*^−^* memory B cells are mostly generated through the germinal center reaction, DN B cells are generated in the interfollicular space where they interact and get activated by T cells, bypassing the germinal center events to undergo affinity maturation and somatic hypermutation (Elsner and Shlomchik, 2020). Such B cells are dominant in several autoimmune conditions such as lupus and in neuro-inflammatory diseases (Jenks et al., 2018; Ruschil et al., 2020). They are also increased early following vaccinations (Ruschil et al., 2020) and are thought to be important for the initial antiviral response that generates short-lived plasmablasts and gradually replaced by germinal center-derived B cell as the infection progresses and the immune system undergoes full activation (Elsner and Shlomchik, 2020; Sokal et al., 2021; Vijay et al., 2020). Persistence of DN B cells could reflect the inability to develop efficient germinal centers as previously reported by us in a more severe form of COVID-19 infection where Bcl-6-expressing T follicular helper cells and germinal centers are not able to form (Kaneko et al., 2020). Alternatively, sustained activation of spike protein-specific B cells in the peripheral reservoir due to viral persistence or antigen spreading could sustain DN B cell development and deregulate the immune response and lead to PCC symptoms. However, our flow cytometry analyses confined to circulating lymphocytes did not permit exhaustive characterization of DN B cell subsets. As the DN subsets are heterogeneous, their detailed characterization and BCR repertoire mapping could provide information on the trajectory of the RBD specific B cell subsets (Jenks et al., 2018; Sanz et al., 2019). Alterations in DN B cell subsets have been observed to be associated with autoantibodies in COVID-19 (Castleman et al., 2022). Even though our analyses of RBD specific DN B cell frequencies did not show significant difference between PCC and No-PCC subjects in both sexes, a possible difference in DN subsets cannot be excluded.

### Coordinated antibody responses

Our observation that the anti-spike and anti-RBD IgG, IgG3 and IgA responses appear to be less coordinated in individuals with PCC points towards the possibility that the propensity to develop PCC may be associated with inherent differences in the evolution of anti-viral immune responses. However, a previous study did not observe differences in the total anti-RBD Ig titers between PCC and No-PCC (Pereira et al., 2021). As the differences between the convalescent groups with and without PCC observed in our study are subtle and nuanced, it is possible that total Ig levels may not truly reflect the differences as each antibody isotype has different functions. For example, Cervia et al. (Cervia et al., 2022) observed that SARS-CoV-2 infected patients with mild or severe disease who went on to develop PCC had higher titers of anti-spike S1 IgG and IgA at the time of infection. Similarly, T cell responses have been observed to be altered in PCC, but the implication of these changes are not clear (Joung et al., 2023; Peluso et al., 2021b; Santa Cruz et al., 2023). Increased antibody responses to the spike antigen following vaccination in the individuals with PCC suggests that differences in immune responses persist over long time after the initial infection is resolved (Joung et al., 2023). These observations suggest that the immune responses to SARS-CoV-2 antigens are qualitatively different in PCC and clearly additional parameters need to be included to predict the immune trajectory toward PCC. In this context, it is noteworthy we observed a strong positive correlation between the number of symptoms at the acute and anti-RBD D614G IgG responses at 1-month post-infection in PCC in males and females.

The human common cold coronaviruses and the three pandemic-associated coronaviruses are not known to cause latent infections (Cui et al., 2019). Infective SARS-CoV-2 viral particles are not known to persist in immunocompetent individuals beyond 9 days after the onset of COVID-19 symptoms even though viral RNA can be detected in respiratory tract, serum and stools for up to 20 days (Cevik et al., 2021; Pan et al., 2021). Nonetheless, both the spike protein and viral RNA has been detected in the circulation of PCC patients (Swank et al., 2022; Tejerina et al., 2022). While we did not detect viral transcripts in the nasal swabs or plasma samples (except for two samples where the amount of virus was below the limit of quantification >100 UI/ml, Ct values of 40.56 and 40.77; data not shown) that were analyzed at 3 months post-infection for viral transcripts, we cannot rule out possible sequestration of viral particles elsewhere as observed in other internal organs (Stein et al., 2022). Fc-gamma receptor-mediated infection of monocytes leads to abortive replication of SARS-CoV-2 and induction of inflammation (Junqueira et al., 2022), suggesting the generation of non-infectious viral particles. ACE2 expression, which mediates viral entry in lung alveolar epithelial cells, has been detected in a wide range of tissues including intestinal enterocytes, vascular endothelium, smooth muscle cells, a subset of dorsal root ganglion neurons, trigeminal sensory ganglia and specialized neuroepithelial cells (Hamming et al., 2004). Despite low levels of viral copy numbers, SARS-CoV-2 has been detected in the brain, kidney, and other organs (Puelles et al., 2020; Stein et al., 2022). Hence, it is possible that viral particles persist in some individuals for a longer duration, possibly due to the infection of various cell types, resulting in prolonged antibody generation contributing to PCC development. Even though coronavirus infections in upper respiratory tract are cleared by humoral and cell mediated immune responses (Callow et al., 1990; Woldemeskel et al., 2020), neuroimaging of PCC patients 10 months after the acute infection points towards certain alterations in functional connectivity and reduction in grey matter in the associated regions (Diez-Cirarda et al., 2022). Using [^18^]F-FDG-PET scan to evaluate glucose uptake by different organs, hypometabolism was observed in the brains of patients with PCC (Guedj et al., 2021a; Guedj et al., 2021b; Sollini et al., 2021). However, it is not known whether PCC is associated with some form of abortive infection of distal organs and tissues in addition to upper respiratory tract. Such a scenario can explain some of the PCC associated symptoms such as the neuro/muscular problems, chronic fatigue and brain fog. It is possible that the heterogeneity in PCC subtypes may be associated with not only the lingering viral particles in the associated organs and tissues but also the variations in immune responses.

### Limitations

One important limitation of our study is the reduced availability of unvaccinated samples after 3 months post infection. This can be explained by the increasing number of people consenting to take the vaccine as recommended by the WHO and national guidelines. The second limitation is the lack of detailed analyses of the various RBD specific B cell subsets including the DN B cells and plasmablasts caused by the access to a very limited quantity of peripheral blood samples. Nonetheless, the results presented in this study support the notion that PCC could be associated with dysregulated immune responses to the viral antigens, at least in males. As we did not analyze germinal center responses, it is not clear whether the observed differences in antibody responses can be considered a proxy for germinal center reactions. Furthermore, whether these altered immune responses are a consequence or a disease promoter awaits further studies in larger and diverse cohorts using additional clinical and immune parameters.

## Acknowledgements

We want to acknowledge Roche Diagnostics Canada for providing the cobas® SARS-CoV-2 Duo kits.

## Author contributions

Study design and funding: SR, AP, IJD, HAC, SI, AC and AM; Identification of clinical samples: CR-P; Sample preparations for flow cytometry, data acquisition and analysis: M-AL and HAC; ELISA: AJIQ, M-AL, FR, SAA, MoN, MaN and MMHC and SR; LP and YD provided reagents; Viral detection in plasma samples: AP, SL and CR-P; Data compilation and analyses: M-AL and SR; PCA analysis: JFL; Statistical analyses: SLP; manuscript writing: M-AL, HAC, AM, SI, AP and SR; Manuscript editing, reviewing: M-AL, AJIQ, SR, AP, IJD, HAC, SI, AC and AM.

## Institutional Review Board Statement

The human studies were approved by the Institutional human ethics committee, CIUSSS de l’Estrie – CHUS, Université de Sherbrooke (approval number 2022-4415).

