## supplemental figure for "SARS-CoV-2 spike antigen-specific B cell and antibody responses in pre-vaccination period COVID-19 convalescent males and females with or without post-covid condition"

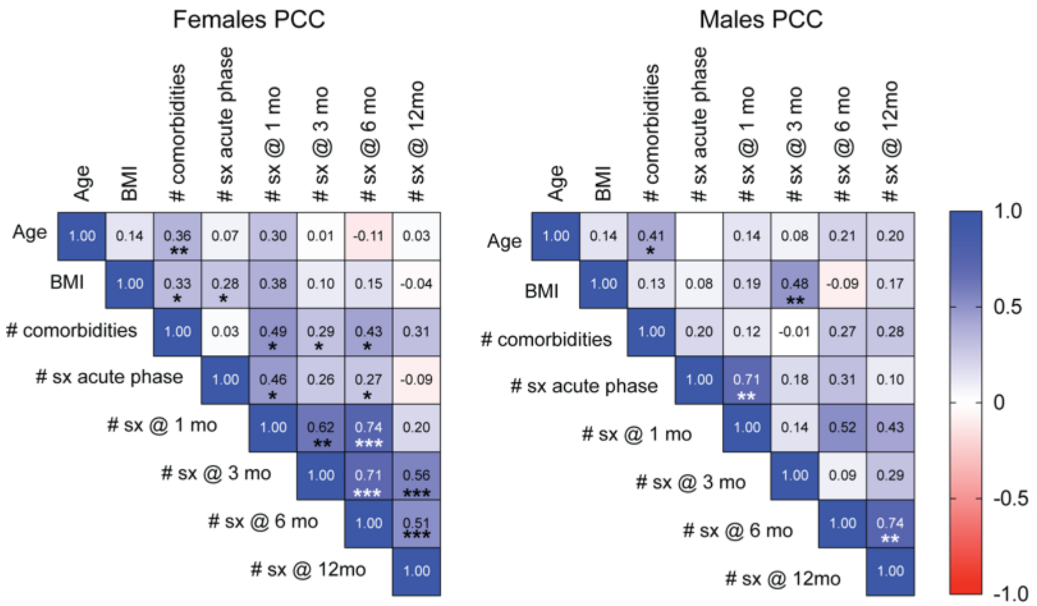


**Supplementary Figure S1. Correlation between clinical parameters of acute disease and PCC in females and males.**

Correlation matrix was generated between age, co-morbidities and the number of symptoms at the acute phase and at 1, 3, 6 or 12-months post-infection in convalescent COVID-19 females and males with PCC. The numbers in the squares indicate the Spearman coefficient value. Asterisks indicate the *p* values: * $p<0.05$; ** $p<0.01$; *** $p<0.001$. Actual *p* values are given in Supplementary Table S4. # sx- number of symptoms at the indicated months post-infection.

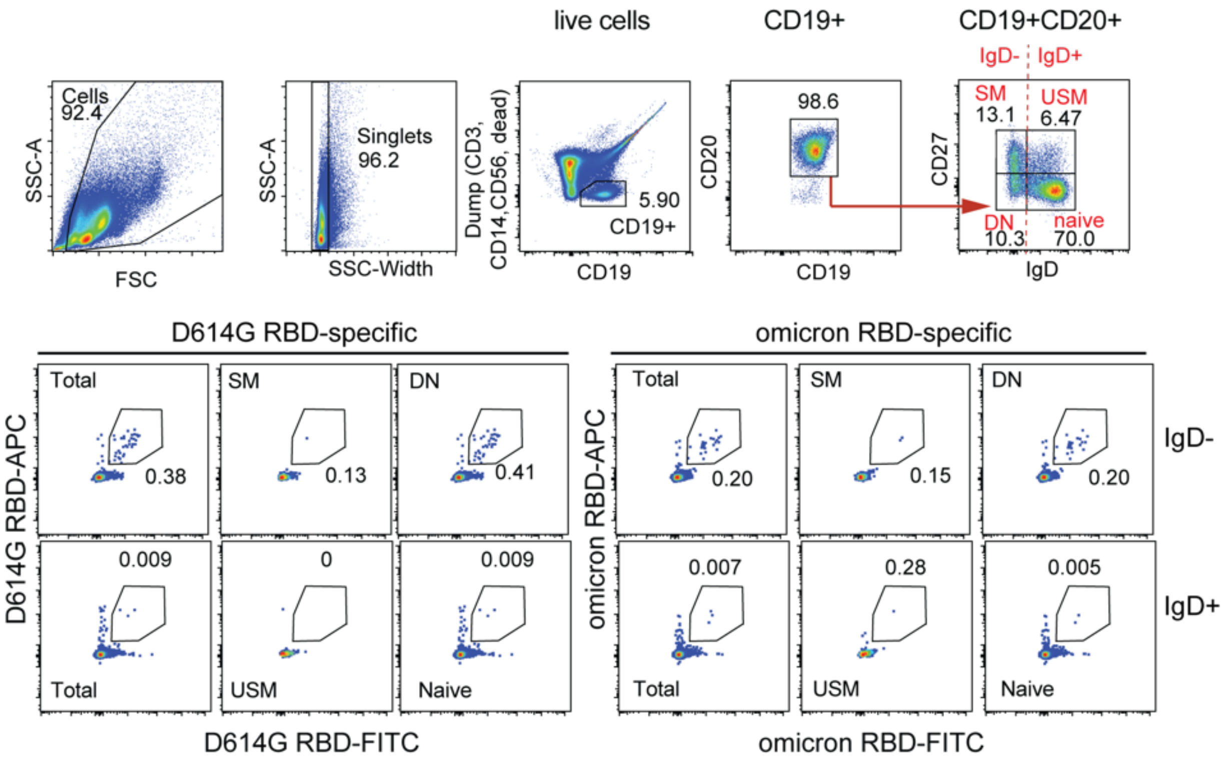


**Supplementary Figure S2. Gating strategy to quantify RBD-specific B cells.**

PBMCs were labelled with B cell markers (CD19, CD20, CD27 and IgD), AF-700 conjugated antibodies for T cells, monocytes and NK (dump channel), D614G-RBD and Omicron-RBD. Data was acquired with Cytoflex-30 cytometer and analyzed by FlowJo v10.3. a) $CD19^{+}CD20^{+}$ B lymphocytes were selected from the singlets after dumping T cells, NK cells and monocytes. CD27 and IgD expression within the B cells defined four subpopulations: naïve ($CD27^{-}IgD^{+}$), unswitched memory (USM; $CD27^{+}IgD^{+}$) switched memory (SM; $CD27^{+}IgD^{-}$), and double negative (DN; $CD27^{-}IgD^{-}$) B cells. Within these B cell subpopulations and total B cells the frequencies of D614G-RBD and Omicron-RBD reactive cells were estimated.

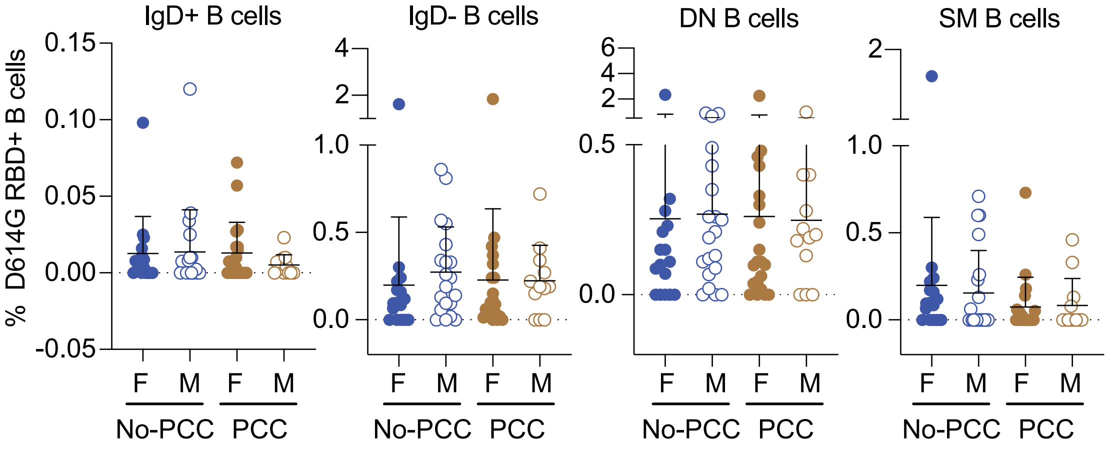


**Supplementary Figure S3: D614G-RBD specific B cell frequencies in convalescent PCC and No-PCC groups at 3 months post-infection.**

Data presented in Fig. 2 were segregated by sex and compared. No significant differences were observed between females and males in the frequencies of D614G RBD reactive B cell subsets as evaluated by Mann Whitney's test.

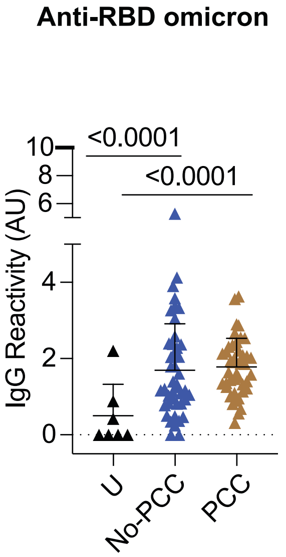


**Supplementary Figure S4. Anti- omicron RBD specific IgG responses in uninfected and convalescent individuals with or without PCC.**

Plasma samples were collected during routine clinical visit at 3 months post PCR-positive diagnosis. Anti-omicron IgG responses in uninfected (U), No-PCC and PCC groups were determined by ELISA. The groups were compared by Mann Whitney's test.

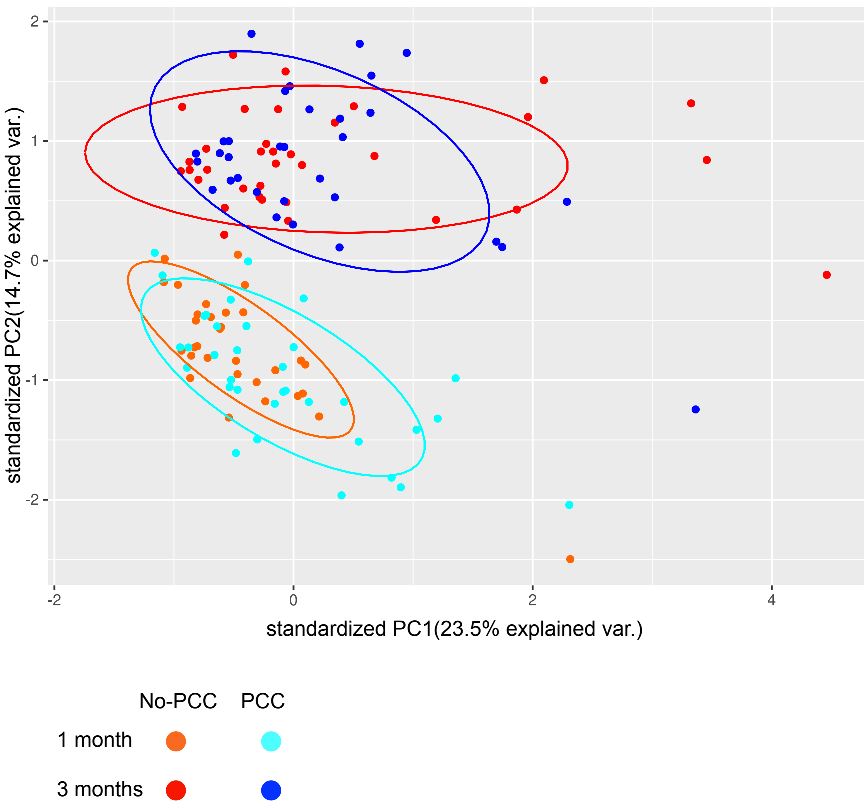


**Supplementary Figure S5. PCA analyses of clinical parameters, B cell reactivities and antibody responses to SARS-CoV-2 antigens in PCC and No-PCC groups at 1 and 3 months post-infection.** The PCA plot was generated using the R 4.1.2 (https://cran.r-project.org/) ggbiplot package (https://rdocumentation.org/packages/ggbiplot/versions/0.55). Immune parameters for which more than 50% of the samples had values at both 1- and 3-month timepoints were utilized. These parameters included sex, number of symptoms at the acute infection timepoint, BMI, anti-D614G RBD specific B cell subsets ($CD19^+$, $CD19^+CD20^+$, $IgD^+$, $IgD^-$, naïve, DN, SM and USM), anti-D614G RBD IgG, anti-spike IgG, anti-spike IgG3, anti-spike IgA, anti-nucleocapsid IgG and anti-nucleocapsid IgA. Excluded from the data were data points with no values as well as those obtained after vaccination. PC1 and PC2 were plotted and a normal data ellipse for each group was generated.

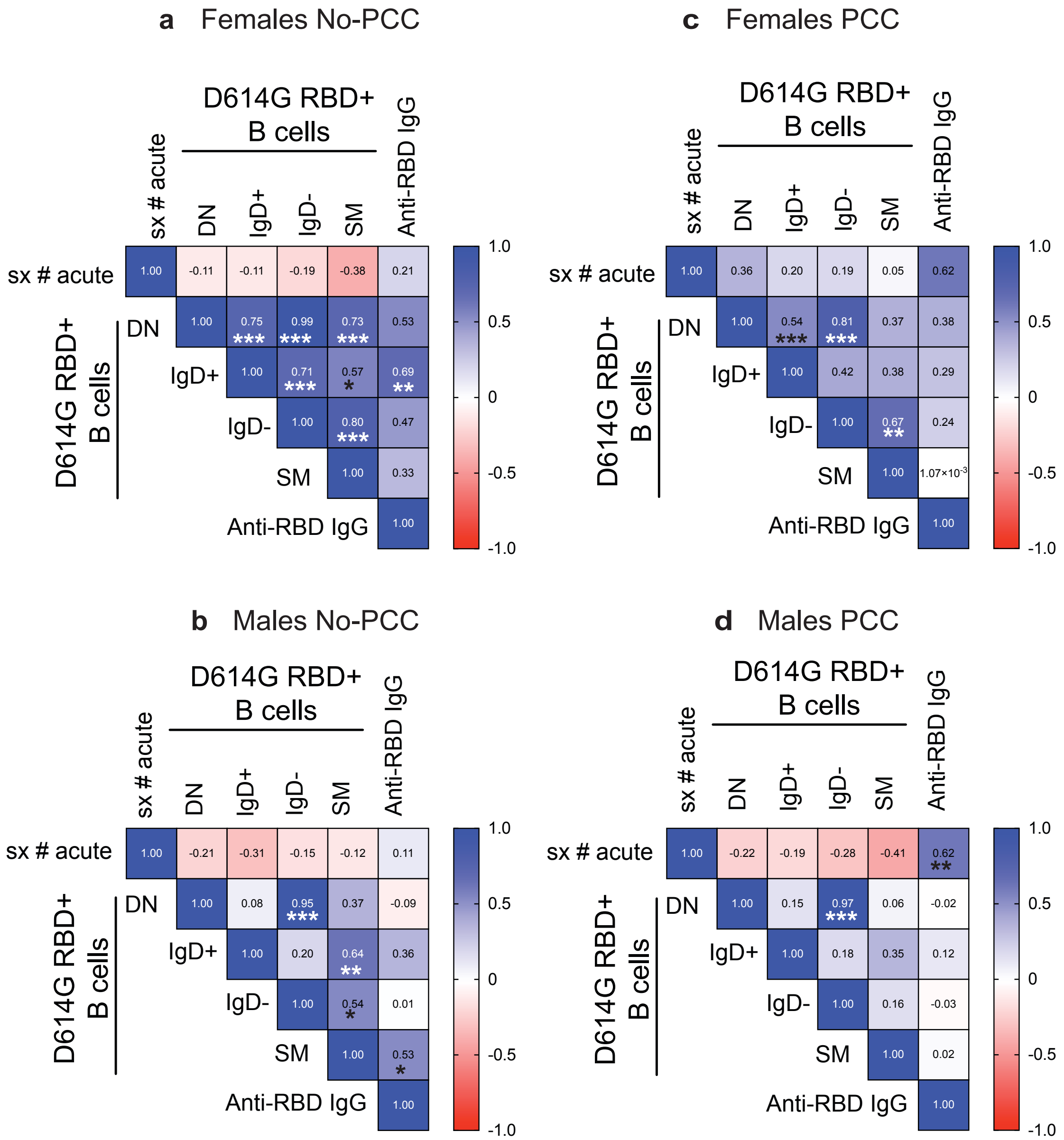


**Supplementary Figure S6.** Correlation matrix for B cell responses at 1- month post infection in females and males with and without PCC. The numbers in the squares indicate the Spearman coefficient value. The *p* values are indicated in the figure as asterisks ( * $p$<0.05, ** $p$<0.01, *** $p$<0.001) and are given in **Supplementary Table S6**. For simplicity they are indicated in the figure with asterisks.

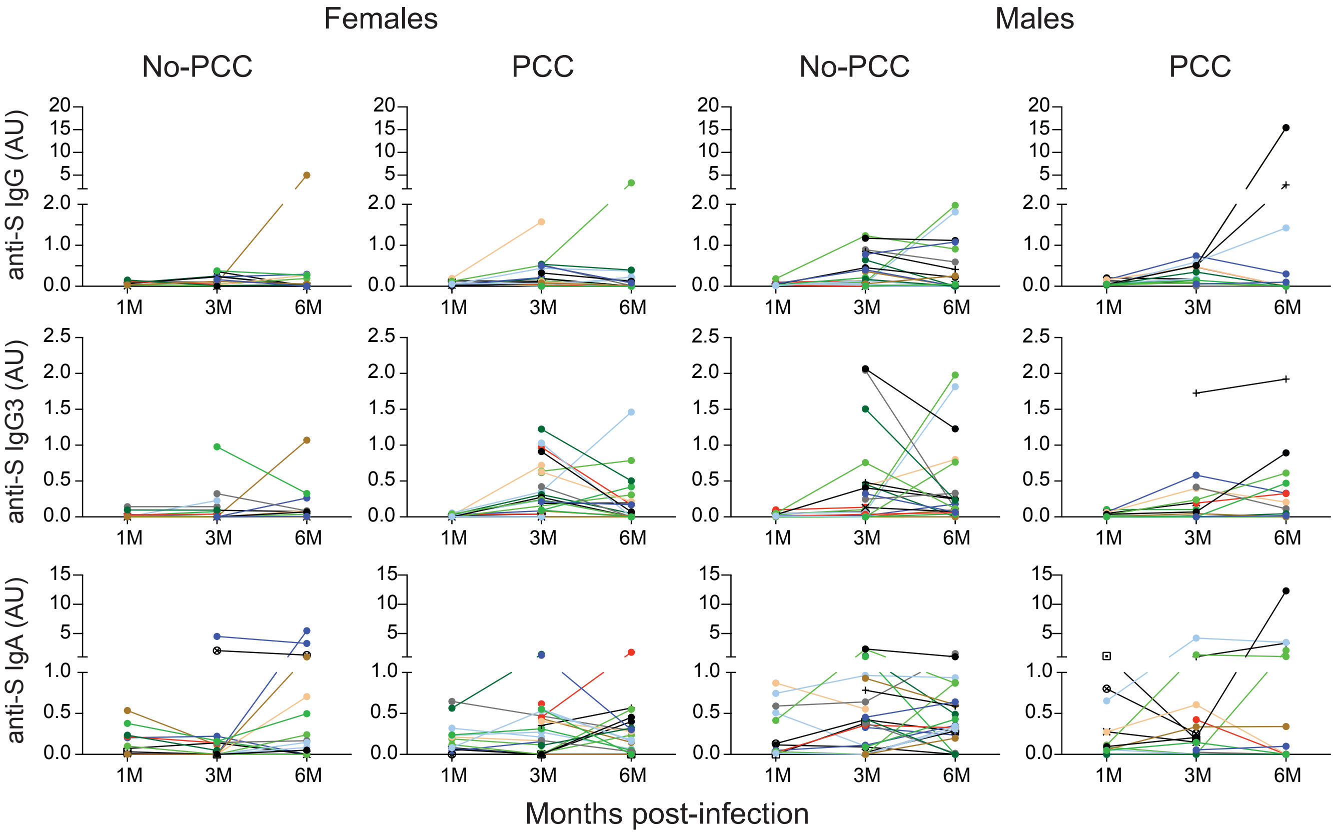


**Supplementary Fig 7. Evolution of anti-spike IgG, IgG3 and IgA antibody response from 1 month to 6 months in females and males with or without PCC.**

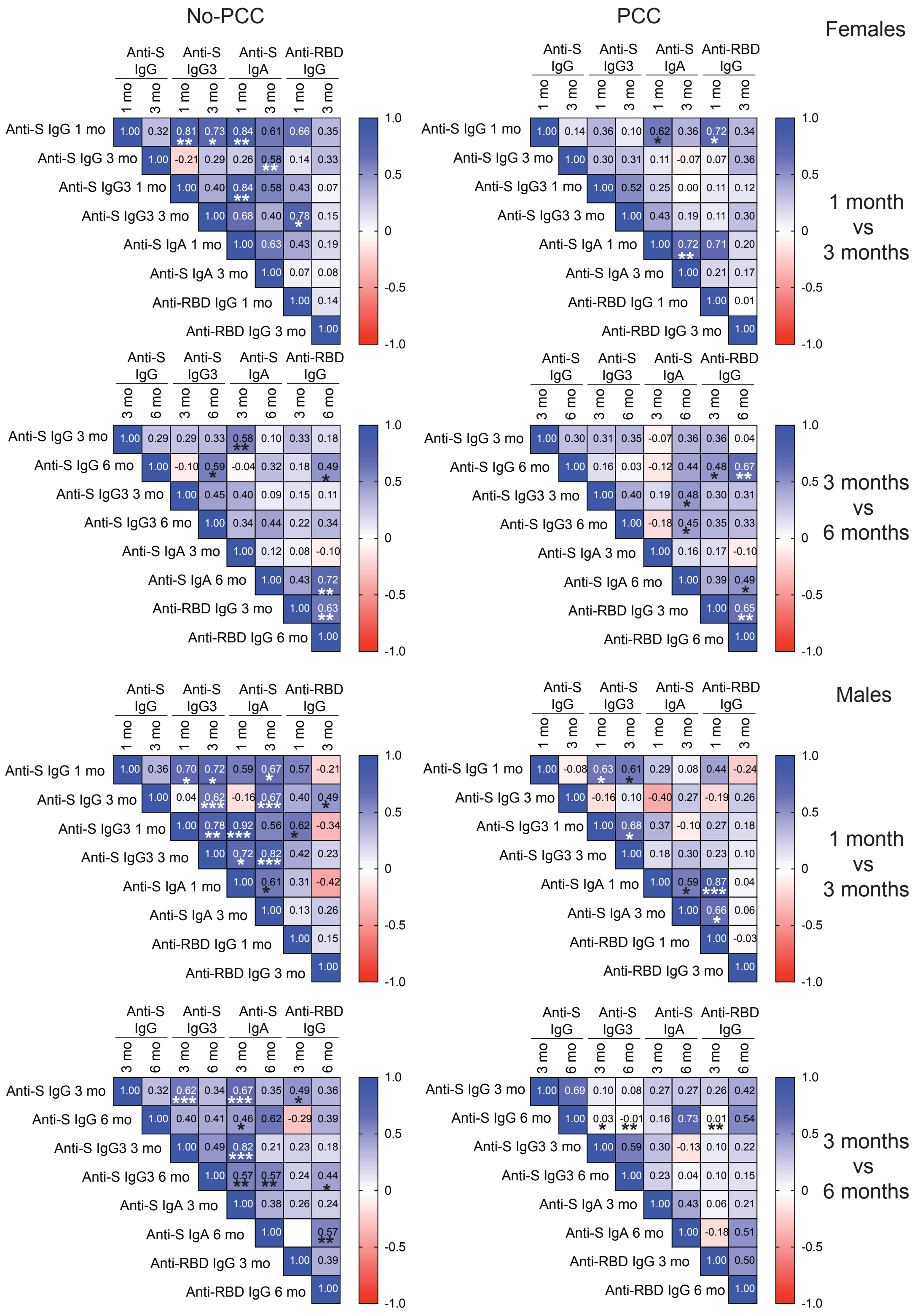


**Supplementary Fig. 8. Correlation matrix for antibody responses between 1- and 3- months and 3- and 6-months post infection in females and males with and without PCC.** RBD refers to RBD-

D614G. The numbers in the squares indicate the Spearman coefficient value. The *p* values are indicated in the figure as asterisks ( * $p<0.05$, ** $p<0.01$, *** $p<0.001$) and are given in **Supplementary Table S7**.

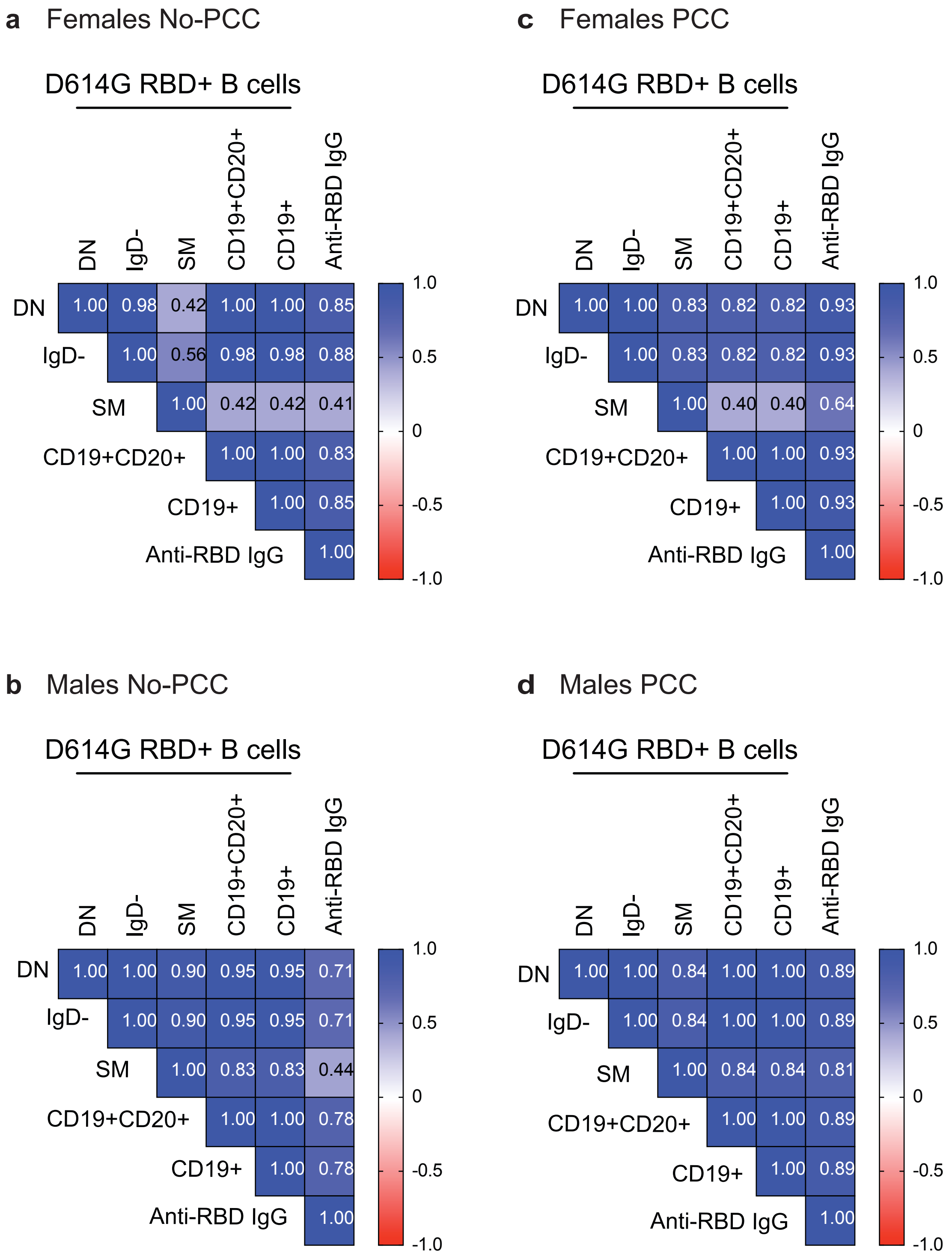

**Supplementary Fig. S9. Correlation matrix for anti-RBD responses at 12 in all the samples irrespective of their vaccination status.** The numbers in the squares indicate the Spearman coefficient value. The *p* values given in **Supplementary Table S8** are not significant**.**

**Supplementary Table S1.** Antibodies and reagents used for flow cytometry.

| Reagent | Source | Cat. # | Clone |
|---|---|---|---|
| Anti-human CD3 - AF700 | BioLegend | 300424 | UCHT1 |
| Anti-human CD14 - AF700 | BioLegend | 367114 | 63D3 |
| Anti-human CD56 - AF700 | BioLegend | 318316 | HCD56 |
| Anti-human CD19 - APC-Cy7 | BioLegend | 302218 | HIB19 |
| Anti-human CD27 - BV605 | BioLegend | 302830 | O323 |
| Anti-human CD20 - PE-Cy7 | BioLegend | 302312 | 2H7 |
| Anti-human IgD - BV785 | BioLegend | 348242 | IA6-2 |
| RBD D614G* -biotin | AcroBiosystems | SPD-C82E8 | |
| RBD Omicron -biotin | AcroBiosystems | SPD-C82E4 | |
| Streptavidin - APC | BioLegend | 405207 | |
| Streptavidin - FITC | BioLegend | 405202 | |
| DRAQ7 live/dead stain | BioLegend | 424001 | |

* RBD D614G refers to the strain that was in circulation in 2020.

**Supplementary Table S2.** Materials used for ELISA

| Material | Source | Cat # |
|---|---|---|
| 96-well high-binding microtiter plate | Greiner Bio-One | 655061 |
| RBD D614G (ELISA) | AcroBiosystems | SPD-C52H3 |
| RBD Omicron (ELISA) | AcroBiosystems | SPD-C522e |
| Spike Protein | NRC | SMT1-1 |
| Nucleocapsid Protein | NRC | NCAP-1 |
| Anti-human IgG | Sigma | B3773 |
| Anti-human IgA | Sigma | SAB3701227 |
| Anti-human IgG3 | Sigma | B3523 |
| Streptavidine-Peroxidase | Sigma | 18-152 |
| HRP-IgA Conjugate (for N protein) | Sigma | SAB3701236 |
| HRP-IgG Conjugate (for N protein) | Sigma | A0170 |
| TMB | Sigma | ES022 |

**Supplementary Table S3.** Plasma dilutions used for ELISA

| Antibody specificity | Plasma dilution |
|---|---|
| Anti-D614G IgG RBD | 1:1000 |
| Anti-Omi IgG RBD | 1:500 |
| Anti-Spike IgG | 1:1000 |
| Anti-Spike, RBD IgG3 | 1:100 |
| Anti-Spike, RBD IgA | 1:500 |
| Anti-N IgG | 1:1000 |
| Anti-N IgA | 1:500 (3 months post-infection) |
| Anti-N IgA | 1:1000 (1 month post-infection) |
